# Disentangling temporal associations in marine microbial networks

**DOI:** 10.1101/2021.07.13.452187

**Authors:** Ina Maria Deutschmann, Anders K. Krabberød, Francisco Latorre, Erwan Delage, Cèlia Marrasé, Vanessa Balagué, Josep M. Gasol, Ramon Massana, Damien Eveillard, Samuel Chaffron, Ramiro Logares

## Abstract

**Background:** Microbial interactions are fundamental for Earth’s ecosystem functioning and biogeochemical cycling. Nevertheless, they are challenging to identify and remain barely known. Omics-based censuses are helpful in predicting microbial interactions through the statistical inference of single (static) association networks. Yet, microbial interactions are dynamic and we have limited knowledge of how they change over time. Here we investigate the dynamics of microbial associations in a 10-year marine time series in the Mediterranean Sea using an approach inferring a time-resolved (temporal) network from a single static network.

**Results:** A single static network including microbial eukaryotes and bacteria was built using metabarcoding data derived from 120 monthly samples. For the decade, we aimed to identify persistent, seasonal, and temporary microbial associations by determining a temporal network that captures the interactome of each individual sample. We found that the temporal network appears to follow an annual cycle, collapsing and reassembling when transiting between colder and warmer waters. We observed higher association repeatability in colder than in warmer months. Only 16 associations could be validated using observations reported in literature, underlining our knowledge gap in marine microbial ecological interactions.

**Conclusions:** Our results indicate that marine microbial associations follow recurrent temporal dynamics in temperate zones, which need to be accounted for to better understand the functioning of the ocean microbiome. The constructed marine temporal network may serve as a resource for testing season-specific microbial interaction hypotheses. The applied approach can be transferred to microbiome studies in other ecosystems.

## INTRODUCTION

Microorganisms are the most abundant life forms on Earth, being fundamental for global ecosystem functioning [1–3]. The total number of microorganisms on the planet is estimated to be ≈ 10^12^ species [4] and ≈ 10^30^ cells [5, 6]. In particular, microorganisms dominate the largest biome, the ocean, which harbors ≈ 10^29^ microbial cells [6] accounting for ~70% of the total marine biomass [7, 8].

Microbial communities are highly dynamic and their composition is determined through a combination of ecological processes: selection, dispersal, drift, and speciation [9]. Selection is a prominent community structuring force exerted via multiple abiotic and biotic factors [10, 11]. Several studies have addressed the role of *abiotic* factors in structuring microbial communities. For example, temperature is one of the main factors exerting selection in the ocean microbiome over spatiotemporal scales [12–15]. *Biotic* factors can also strongly affect microbial communities [16]. However, a mechanistic understanding of how they affect community structure is currently lacking, as the diversity of microbial interactions is barely known [3, 17].

The vast microbial diversity and the fact that most microorganisms are still uncultured [18, 19] make it impossible to experimentally test all potential interactions between pairs of microbes. However, omics-technologies allow estimating microbial relative abundances over spatiotemporal scales, which permits determining statistical associations between taxa. These associations can be summarized as a network with nodes representing microorganisms and edges representing potential interactions [20, 21].

As microorganisms are highly interconnected [21], association networks provide a general overview of the entire microbial system and have been tremendously valuable for generating novel hypotheses about putative interactions. In particular, time series have allowed identifying potential ecological interactions among marine microorganisms [22–28]. For example, previous work characterized ecological links between marine archaea, bacteria, and eukaryotes [22], including links with viruses [24, 26], also investigating within- and between ocean-depth relationships [25, 27]. These studies not only identified time-dependent associations among ecologically important taxa, but also potential synergistic or antagonistic relationships, as well as possible ‘keystone’ species and potential niches [22, 23]. Moreover, several studies have reported more associations among microorganisms than between microorganisms and environmental variables, suggesting the importance of biotic relationships in structuring microbial community assemblages [22, 28].

Previous studies have used temporal microbial abundance data to infer static networks summarizing all potential associations in space and time. This static abstraction assumes that the network topology does not change (static) and edges represent persistent associations assumed as interactions [29]; that is, edges are present throughout time and space. This assumption cannot represent the reality of most microbial interactions. Thus, a single static network usually contains persistent, temporary, and recurring (including seasonal) associations that need to be disentangled.

Despite the contribution of static networks to our understanding of microbial interactions in the ocean, it is necessary to incorporate the temporal dimension. Using a time-resolved, i.e., temporal network instead of a single static network would allow investigation of the dynamic nature of microbial associations and how they change over time, whether the change is deterministic or stochastic, and how environmental selection influences network architecture. Addressing these questions is fundamental for a better understanding of the dynamic interactions that underpin ecosystem function in the ocean. Here, we investigated marine microbial associations through time by determining a temporal network from a single static network.

## MATERIALS AND METHODS

### The Blanes Bay Microbial Observatory (BBMO)

The BBMO is a coastal oligotrophic site in the North-Western Mediterranean Sea (41*°*40’N 2*°*48’E) without major natural disturbances and little anthropogenic pressure, except for the construction of a nearby harbor between 2010 and 2012 [30, 31]. The seasonal cycle is typical for a temperate coastal system [30], and the main environmental factors influencing seasonal microbial succession have been well studied and are known [12]. Shortly, the water column is slightly stratified in summer before it destabilizes and mixes with water from offshore in late fall, increasing the availability of inorganic nutrients with maximum concentrations in winter, between November and March. The high amount of nutrients and increasing light induce phytoplankton blooms, mostly in late winter-early spring. During summer, inorganic nutrients become limiting, the primary production is minimal, and dissolved organic carbon accumulates [30].

### From sampling to microbial relative abundances

We sampled surface water (*≈* 1m depth) monthly from January 2004 to December 2013 to determine microbial community composition and also measured ten environmental variables, which were previously described [13, 30]: water temperature (°C) and salinity (obtained *in situ* with a SAIV-AS-SD204 Conductivity-Temperature-Depth probe), day-length (hours of light), turbidity (Secchi depth in meters), total chlorophyll-a concentration (*μ*g/l, fluorometry of acetone extracts after 150 ml filtration on GF/F filters [30]), and five inorganic nutrients: PO_4_^3-^, NH_4_^+^, NO_2_^-^, NO_3_^-^ and SiO_2_ (*μ*M, determined with an Alliance Evolution II autoanalyzer [32]).

Sampling of microbial communities, DNA extraction, rRNA-gene amplification, sequencing, and bioinformatic analyses are explained in detail in [28]. In short, 6 L of water were prefiltered through a 200 μm nylon mesh and subsequently filtered through another 20 μm nylon mesh and separated into nanoplankton (3 – 20 μm) and picoplankton (0.2 – 3 μm) using a 3 μm and 0.2 μm pore-size polycarbonate and Sterivex filters, respectively. Then, the DNA was extracted from the filters using a phenol-chloroform protocol [33], which has been modified for purification with Amicon units (Millipore). We amplified the 18S rRNA genes (V4 region) with the primers TAReukFWD1 and TAReukREV3 [34], and the 16S rRNA genes (V4 region) with Bakt 341F [35] and 806RB [36]. Amplicons were sequenced in a MiSeq platform (2×250bp) at RTL Genomics (Lubbock, Texas). Read quality control, trimming, and inference of Operational Taxonomic Units (OTUs) delineated as Amplicon Sequence Variants (ASVs) were done with DADA2 [37], v1.10.1, with the maximum number of expected errors set to 2 and 4 for the forward and reverse reads, respectively.

Microbial sequence abundance tables were obtained for each size fraction for both microbial eukaryotes and prokaryotes. Before merging the tables, we subsampled each table to the lowest sequencing depth of 4907 reads with the *rrarefy* function from the Vegan R-package [38], v2.4-2, (see details in [28]). We excluded 29 nanoplankton samples (March 2004, February 2005, May 2010 - July 2012) due to suboptimal amplicon sequencing. In these samples, abundances were estimated using seasonally aware missing value imputation by the weighted moving average for time series as implemented in the *imputeTS* R-package, v2.8 [39]. These imputed values did not introduce biases in the analyses [28].

Sequence taxonomy was inferred using the naïve Bayesian classifier method [40] together with the SILVA database [41], v.132, as implemented in DADA2 [37]. Additionally, eukaryotic microorganisms were BLASTed [42] against the Protist Ribosomal Reference (PR2) database [43], v4.10.0. The PR2 classification was used when the taxonomic assignment from SILVA and PR2 disagreed. We removed ASVs that were identified as Metazoa, Streptophyta, plastids, mitochondria, and Archaea since the 341F-primer is not optimal for recovering this domain [44]. Besides, Haptophyta is known to be missed by the primer TAReukREV3 [45].

The resulting table contained 2924 ASVs (Table 1A). Next, we removed rare ASVs keeping ASVs with sequence abundance sums above 100 reads and prevalence above 15% of the samples, i.e., we considered taxa present in at least 19 months. The resulting table contained 1782 ASVs (Table 1B). An ASV can appear twice, in the nanoplankton and picoplankton size fractions. However, an ASV may be detected in both size fractions due to dislodging cells or particles and filter clogging, which can introduce biases in our analysis. To reduce these biases, and as done previously [28], we divided the abundance sum of the larger by the smaller size fraction for each ASV appearing in both size fractions and set the picoplankton abundances to zero if the ratio exceeded 2. Likewise, we set the nanoplankton abundances to zero if the ratio was below 0.5. This operation removed two eukaryotic ASVs and 41 bacterial ASVs from the nanoplankton, and 30 bacterial ASVs from the picoplankton (Table 1C). The resulting abundance table was used for network inference.

**Table 1:** Number and fraction of ASVs and reads (total, bacterial and eukaryotic) for the sequence abundance tables (A, B, and C), the preliminary network with significant edges (D), and the single static network (E) obtained after removing environmentally-driven edges and edges with association partners appearing more often alone than with the partner. If an ASV appeared in the nano- and pico-plankton size fractions, it was counted twice.

| Count tables | ASVs | Reads | Eukaryote | Eukaryotic reads | Bacteria | Bacterial reads |
| --- | --- | --- | --- | --- | --- | --- |
| A | 2 924 | 2 273 548 | 1 365 | 1 121 855 | 1 559 | 1 151 693 |
| B | 1 782 | 2 155 318 | 1 009 | 1 057 599 | 773 | 1 097 719 |
| C | 1 709 | 2 062 866 | 1 007 | 1 057 263 | 702 | 1 005 603 |
| D | 754 | 1 657 885 | 306 | 730 025 | 448 | 927 860 |
| E | 709 | 1 621 959 | 294 | 719 558 | 415 | 902 401 |

| Fractions | ASV | Reads | Eukaryote | Eukaryotic reads | Bacteria | Bacterial reads |
| --- | --- | --- | --- | --- | --- | --- |
| B/A*100 | 60.94 | 94.80 | 73.92 | 94.27 | 49.58 | 95.31 |
| C/A*100 | 58.45 | 90.73 | 73.77 | 94.24 | 45.03 | 87.32 |
| D/C*100 | 44.12 | 80.37 | 30.39 | 69.05 | 69.05 | 92.27 |
| E/C*100 | 41.49 | 78.63 | 29.20 | 68.06 | 59.12 | 89.74 |

*A – raw sequence abundance table; B – sequence abundance table without rare ASVs; C – sequence abundance table after size-fraction filtering; D – preliminary network with significant edges; E – single static network*

### From sequence abundances to the single static network

First, we constructed a preliminary network using the tool eLSA [46, 47], as done in [28, 48], including default normalization and z-score transformation, using median and median absolute deviation. Although we are aware of time-delayed interactions, we considered our 1-month sampling interval too large for inferring time-delayed associations with a solid ecological basis, and focused on contemporary interactions between co-occurring microorganisms. Using 2000 iterations, we estimated p-values with a mixed approach that performs a random permutation test of a co-occurrence if the comparison’s theoretical *p*-values are below 0.05. The Bonferroni false discovery rate (*q*) was calculated based on the *p*-values using the *p.adjust* function from the stats R-package [49]. We used the 0.001 significance threshold for the *p* and *q* values, as suggested in other studies [20]. We refrained from using an association strength threshold since it may not be appropriate to differentiate between true interactions and environmentally-driven associations [48]. Furthermore, changing thresholds have been shown to lead to different network properties [50]. The preliminary network contained 754 nodes and 29820 edges (24458, 82% positive, and 5362, 18% negative).

Second, for environmentally-driven edge detection, we applied EnDED [48], combining the methods Interaction Information (with a 0.05 significance threshold and 10000 iterations) and Data Processing Inequality. We inserted artificial edges connecting each node to each environmental parameter. We identified and removed 3315 (11.12%) edges that were environmentally driven; 26505 edges (23405, 88.3% positive, and 3100, 11.7% negative) remained (Supplementary Tables 3 and 4).

Third, we determined the Jaccard index, *J*, for each microorganisms pair associated through an edge, in order to remove associations between microorganisms that have a low co-occurrence. Let *S_i_*, be the set of samples in which both microorganisms are present (sequence abundance above zero), and *S_u_* be the set of samples in which one or both microorganisms are present. Then, we can calculate the Jaccard index as the fraction of samples in which both appear (intersection) from the number of samples in which at least one appears (union): *J* = *S_i_*/*S_u_*. We chose *J* > 0.5 as in previous work [48], which removed 9879 edges and kept 16626 edges (16481, 99.1% positive and 145, 0.9% negative). We removed isolated nodes, i.e., nodes without an associated partner in the network. The number and fraction of retained reads are listed in Table 1. The resulting network is our single static network.

### From the single static network to the temporal network

We determined the temporal network comprising 120 sample-specific (monthly) subnetworks through the three conditions indicated below and visualized in Figure 1. The subnetworks are derived from the single static network and contain a node subset and an edge subset of the static network. Let *e* be an association between microorganisms *A* and *B*, with association duration *d* = (*t*_1_, *t*_2_), i.e., the association starts at time point *t*_1_ and ends at *t*_2_. Then, considering month *m*, the association *e* is present in the monthly subnetwork *N_m_*, if

1. *e* is an association in the single static network,
2. the microorganisms *A* and *B* are present within month *m*, and
3. *m* is within the duration of association, i.e., *t*_1_ ≤ *m* ≤ *t*_2_.

**Figure 1:**
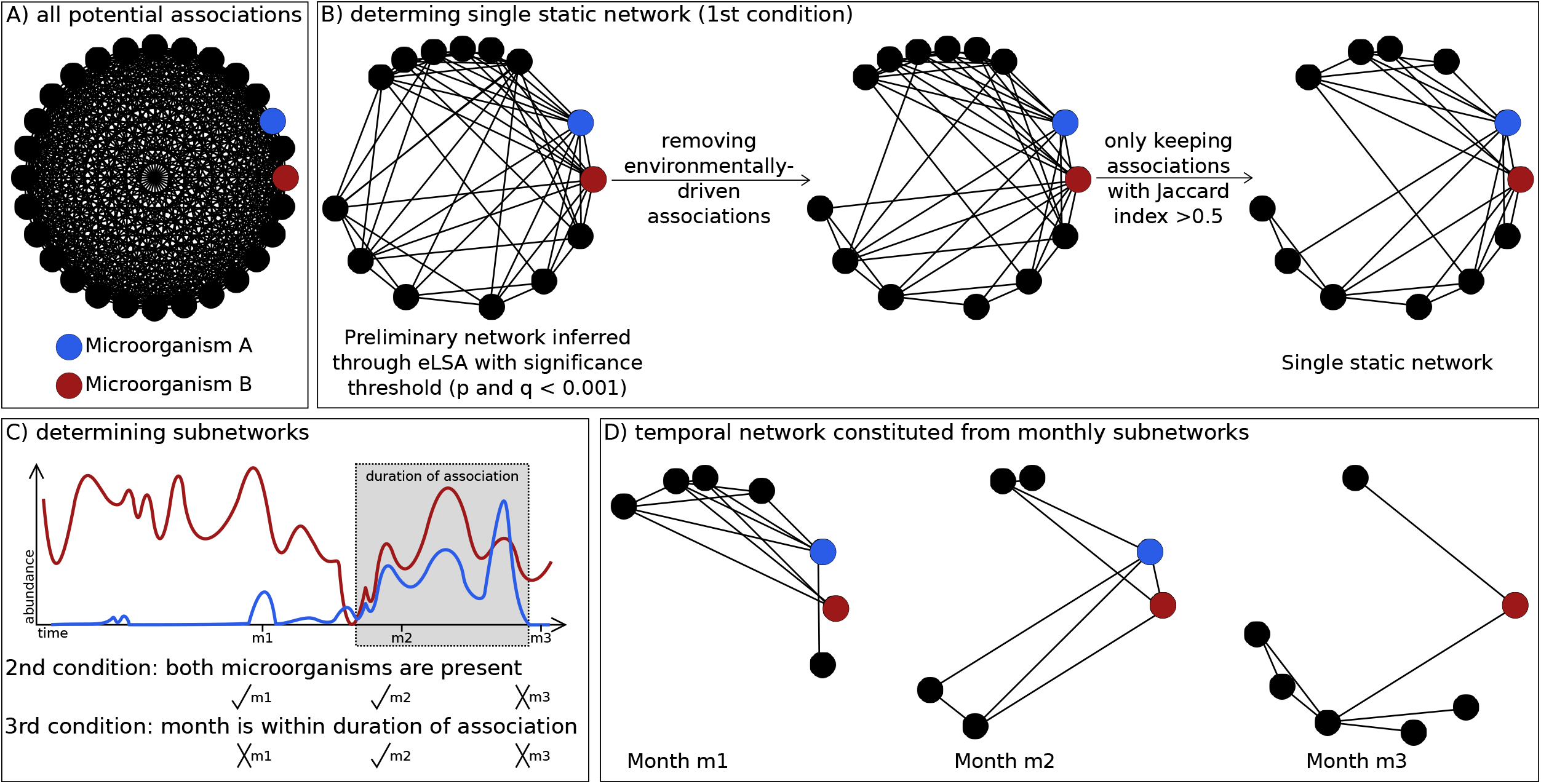
Estimating a temporal network from a single static network via subnetworks. A) A complete network would contain all possible associations (edges) between microorganism (nodes). B) The single static network inferred with the network construction tool eLSA and the applied filtering strategy considering association significance, the removal of environmentally-driven associations, and associations whose partners appeared in more samples together than alone, i.e., Jaccard index being above 0.5. An association having to be present in the single static network is the first out of the three conditions for an association to be present in a monthly subnetwork. C) In order to determine monthly subnetworks, we established two further conditions for each edge. First, both microorganisms need to be present in the sample taken in the specific month. Second, the month lays within the time window of the association inferred through the network construction tool. Here, three months are indicated as an example. D) Example of monthly subnetworks for the three months. The colored nodes correspond to the abundances depicted in C).

With the 2^nd^ condition, we assumed that an association was present in a month if both microorganisms were present, i.e., the microbial abundances were non-zero for that month. However, we cannot assume that microbial co-occurrence is a sufficient condition for a microbial interaction because different mechanisms influence species and interactions, and the environmental filtering of species and interactions can differ [51]. Using only the species occurrence assumption would increase association prevalence. To lower this bias, we also required that the association was present in the static network, 1^st^ condition, and within the association duration, 3^rd^ condition, both inferred by eLSA [46, 47]. Lastly, we removed isolated nodes from each monthly subnetwork.

### Network analysis

We computed global network metrics to characterize the single static network and each monthly subnetwork using the igraph R-package [52]. Some metrics tend to be more correlated than others implying redundancy between them, grouping them into four groups [53]. Thus, we selected one metric from each group: *edge density, average path length, transitivity,* and *assortativity* based on node degree. In addition, we also computed the *average strength of positive associations* between microorganisms using the mean, and *assortativity* based on the nominal classification of nodes into bacteria and eukaryotes. Assortativity (bacteria vs. eukaryotes) is positive if bacteria tend to connect with bacteria and eukaryotes tend to connect with eukaryotes. It is negative if bacteria tend to connect to eukaryotes and vice versa. We also quantified associations by calculating their prevalence as the fraction of monthly subnetworks in which the association was present for all ten years (recurrence), and monthly. We visualized highly prevalent associations with the *circlize* R-package [54]. We tested our hypotheses of environmental factors influencing network topology by calculating the Spearman correlations between global network metrics and environmental data. We used Holm’s multiple test correction to adjust p-values [55], with the function *corr.test* in the *psych* R-package [56]. We used Gephi [57], v.0.9.2, and the Fruchterman Reingold Layout [58] for network visualizations.

### Test of network construction tool

We have used eLSA to estimate the duration of an association, which we used as the third condition (*m* is within the duration of association, i.e., *t*_1_ ≤ *m* ≤*t*_2_) to infer the sample-specific subnetworks. Other methods may perform better on compositional data such as ours [59] (although this is not necessarily the case; see [60]). Therefore we tested another network construction approach (FlashWeave [61]) for comparative purposes. FlashWeave performed better than eLSA in some benchmark tests run by other authors, while eLSA performed better than FlashWeave in other tests [61]. FlashWeave can handle sparse datasets taking zeros into account and avoiding spurious correlations between ASVs that share many zeros. However, it neglects the temporal variation. To control data compositionality [59], we applied a centered-log-ratio transformation separately to the bacterial and eukaryotic read abundance tables before merging them. Then, we inferred a network using FlashWeave [61], selecting the options “heterogeneous” and “sensitive”. We have run analyses including the environmental data (10 variables; see above). The resulting network had 932 nodes and 1440 edges. Next, we determined a temporal network using conditions 1) and 2) but not 3) since the temporal duration is not estimated by FlashWeave. FlashWeave results are used hereafter to compare against eLSA, although eLSA is kept as the main network construction tool in our work, given that it allows determination of the duration of the associations and there is no evidence suggesting a poor performance of this tool. Thus, unless specified otherwise, we refer to the static and temporal network determined by eLSA.

### Cyanobacteria

Our dataset contained 19 cyanobacterial ASVs, which all appeared in the nano-, and nine in the picoplankton. We blasted the sequences against the Cyanorak database [62], v.2. against the nucleotide database containing all *Synechococcus* and *Prochlorococcus* RNAs with the option -evalue 1.0e-5. We found 2812 sequences comprising 95 different ecotypes (considering name, clade and subclade), with 93.84-100% identity. A total of 11 BBMO ASVs obtained 63 hits with 100% identity, and within these 63 reference sequences there were 34 different ecotypes. Most matching sequences were found for *Synechococcus* ASV_1. While *Synechococcus* ASV_5 had only two 100% hits, they did not 100% match ASV_1 (Supplementary Table 5). Finding *Synechococcus* in both size fractions was against expectations, as this genus is part of the pico-plankton. Yet, they have been observed in fractions above 3 μm at BBMO [63]. Recovering *Synechococcus* ASVs from the nanoplankton may be due to cell aggregation, particle attachment, clogging of filters, or being prey to larger microorganisms. *Synechococcus* could be also picked up in the 3 μm filters during cell division.

### Validated associations

As a general rule, the validation of associations tends to be limited as both true interactions and microorganisms that do not interact with each other are poorly known. As done in [48], we determined true genus-genus interactions as those known in the literature, which are compiled within the Protist Interaction Database, PIDA [17]. On October 15th 2019, PIDA contained 2448 interactions. Although our dataset contains protists and bacteria, we could not evaluate interactions between them through PIDA. The ambiguity in taxonomic classification and the large number of edges challenged the validation. We validated associations between microbial eukaryotes via exact string matching as done previously [48].

## RESULTS

### Extracting a temporal network from a single static association network

From ten years of monthly samples from the Blanes Bay Microbial Observatory (BBMO) in the Mediterranean Sea [30], we computed sequence abundances for 488 bacteria and 1005 microbial eukaryotes from two organismal size-fractions: picoplankton (0.2 – 3 μm) and nanoplankton (3 – 20 μm). We removed Archaea since they are not very abundant in the BBMO surface and primers were not optimal to quantify them. We inferred Amplicon Sequence Variants (ASVs) using the 16S and 18S rRNA-gene. After filtering the initial ASV table for sequence abundance and shared taxa among size fractions, we kept 285 and 417 bacterial and 526 and 481 eukaryotic ASVs in the pico- and nanoplankton size fractions, respectively. We found 214 bacterial ASVs that appeared in both size fractions, but only two eukaryotic ASVs: a *Cryothecomonas* (Cercozoa) and a dinoflagellate (Alveolate).

We used 1709 ASVs to infer a preliminary association network with the tool eLSA [46, 47]. Next, we removed environmentally-driven edges with EnDED [48]. We only considered edges involving partners that co-occurred more than half of the times together than alone (see Methods and Figure 1A-B). Our filtering strategy removed a higher fraction of negative than positive edges (see Methods and Supplementary Table 1). The resulting network is our single static network connecting 709 nodes via 16626 edges (16481 edges, 99.1%, positive and 145, 0.9% negative).

Next, we developed an approach to determine a temporal network. Building upon the single static network, we determined 120 sample-specific (monthly) subnetworks (see Methods for details). These monthly subnetworks represent the 120 months of the time series and together comprise the temporal network. Each monthly subnetwork contains a subset of the nodes and a subset of the edges of the single static network. We used the ASV abundances indicating the presence (ASV abundance > 0) or absence (ASV abundance = 0) as well as the estimated start and duration of associations inferred with the network construction tool eLSA [46, 47] for determining which nodes and edges are present each month (Figure 1, see Methods).

### The single static network metrics differed from most monthly subnetworks

Since each monthly subnetwork was derived from the single static network, they were smaller, containing between 141 (August 2005) and 571 (January 2012) nodes, median ≈354 (Figure 2A), and between 560 (April 2006) to 15704 (January 2012) edges, median ≈6052 (Figure 2B). For further characterization, we computed six global network metrics (Figure 2C and Methods). The results indicated that the single static network differed from most monthly subnetworks and it also differed from the average. In general, the single static network was less connected (edge density) and more clustered (transitivity) with higher distances between nodes (average path length) and stronger associations (average positive association score) than most monthly subnetworks (Figure 2C). In addition, the single static network was usually more assortative according to the node degree but less assortative according to the domain (bacteria vs. eukaryote) than most monthly subnetworks (Figure 2C). High assortativity indicates that nodes tend to connect to nodes of a similar degree and domain.

**Figure 2:**
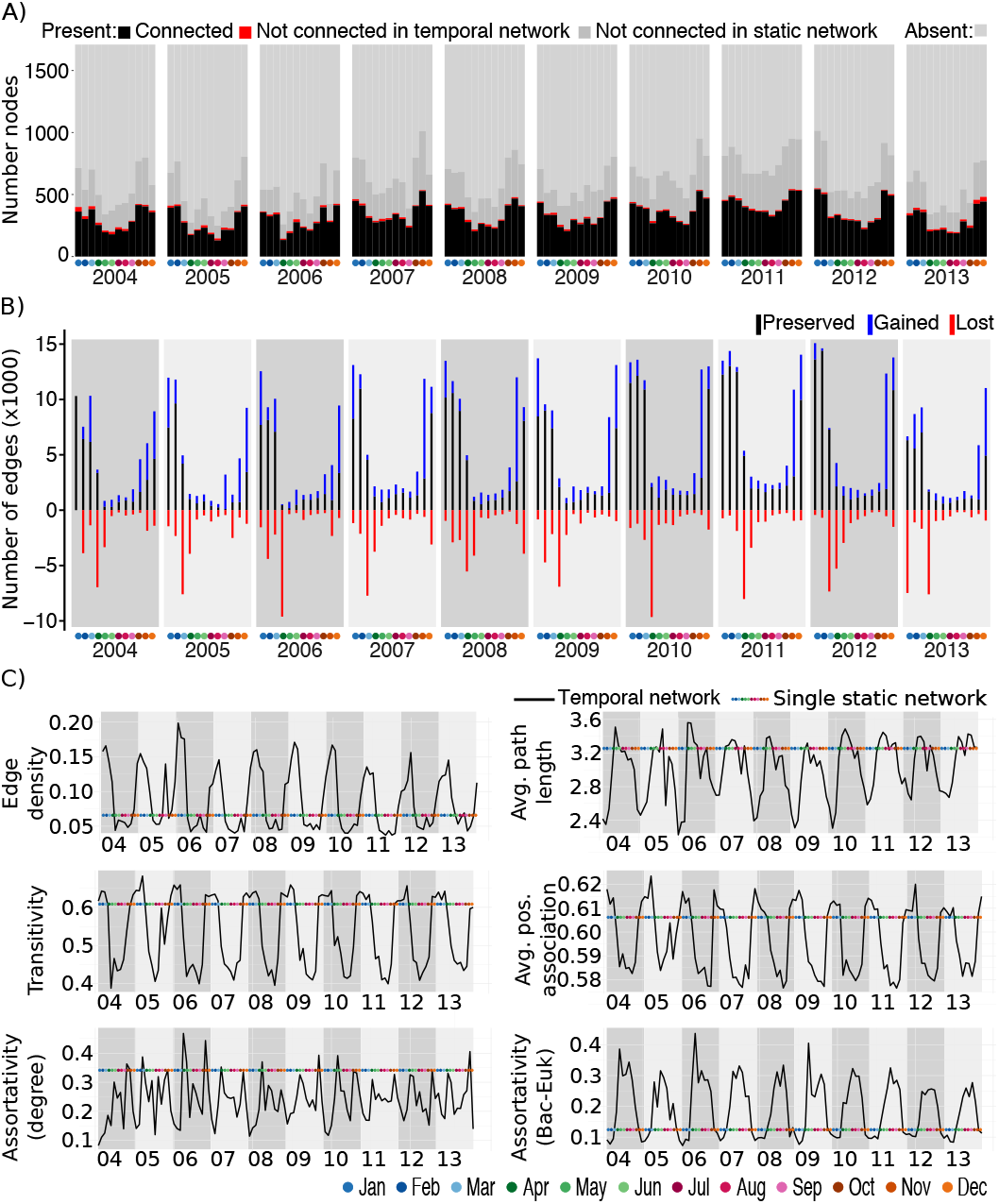
Global (sub)network metrics. A) Number of ASVs (counting an ASV twice if it appears in both size fractions) for each of the 120 months of the Blanes Bay Microbial Observatory time series. There are 1709 ASVs, of which 709 ASVs are connected in the static network. In black, we show the number of nodes connected in the temporal network, and in red the number of nodes that are isolated in the temporal network, i.e., they are connected in the static network and have a sequence abundance above zero for that month (“non-zero”). In dark grey, we show the number of ASVs that are non-zero in a given month but were not connected in the static and subsequently temporal network. In light grey, we show the number of ASVs with zero-abundance in a given month. The sum of connected and isolated nodes and non-zero ASVs represents each month’s richness (i.e., number of ASVs). B) By comparing the edges of two consecutive months, i.e., two consecutive monthly subnetworks, we indicate the number of edges that have been lost (red), preserved (black), and those that are gained (blue), compared to the previous month. C) Six selected global network metrics for each sample-specific subnetwork of the temporal network. The colored line indicates the corresponding metric for the static network.

### Monthly subnetworks display seasonal behavior with yearly periodicity

Over the analyzed decade, the network became more connected and clustered in colder months, with stronger associations and shorter distances between nodes (Figure 2C, Supplementary Figures 1 and 2). Most global network metrics indicated seasonal behavior with yearly periodicity (Figure 2C). For instance, edge density, average positive association score, and transitivity were highest at the beginning and end of each year, while average path length and assortativity (bacteria vs. eukaryotes) were highest in the middle of each year. Assortativity (degree), in contrast to other metrics, usually had two peaks per year corresponding to April-May, and November (Figure 2C). Some metrics (number of nodes and edges, and average path length) presented similar seasonal behavior with yearly periodicity in the temporal network determined from the single static FlashWeave network (Supplementary Figure 3). However, edge density and transitivity displayed patterns contrary to those observed in the temporal network determined from the stingle static eLSA network.

We found mainly temperature and day length, and to a lesser extent nutrient concentrations (mainly SiO_2_, NO_3_^-^ and NO_2_^-^, less PO_4_^3-^), and total chlorophyll-a concentration to affect network topologies as indicated by correlation analyses (Supplementary Figure 2). For example, edge density was highest and temperature lowest in January-March. Then, edge density dropped as temperature increased. April-June displayed edge densities slightly above or similar to those in the warmest months July-September, while October-December had similar or slightly lower edge densities than the coldest months January-March. Edge density vs. hours of light (day length) indicated a yearly recurrent circular pattern for September-April (Supplementary Figure 1). Yet, May-August were not part of the circular pattern. May-August had the highest day length and their corresponding networks low edge density (Supplementary Figure 1).

Next, we quantified how many edges were preserved (kept), lost, and gained (new) in consecutive months. We found the highest loss of edges in April, pointing to a network collapse. The overall number of edges (preserved and gained) was lowest during April-September and increased towards the end of each year (Figure 2B). The number of associations changed over time in a yearly recurring pattern with few associations being preserved when transitioning from colder to warmer waters. We observed a steep network change when transiting from colder to warmer months, reflecting a large reorganization. In turn, the network change from warmer to colder months was less abrupt. Thus, network change between cold and warm waters was not symmetrical over the studied decade at BBMO.

We defined summer and winter as in [28] and compared both seasons between consecutive years in terms of preserved, gained, and lost associations and ASVs. We observed higher repeatability in edges (Supplementary Figure 4) and ASVs (results not shown) in colder than in warmer months, indicating higher predictability during low-temperature seasons.

### Potential core associations

A single static network can comprise permanent, seasonal, and temporary associations. By comparing monthly subnetworks, we identified edges that remain (preserved), appear (gained), or disappear (lost) over time (Figure 2B). Intuitively, we would classify permanent associations through 100% recurrence. However, no association fulfilled the 100% criteria. Most associations had a low recurrence, with three-quarters of the associations present in no more than 38% (total 46) of the monthly subnetworks. The average association prevalence was similar across taxonomic ranks (Supplementary Figure 5). Considering the 100 most prevalent associations, which appeared in 71.7-98.3% (total 86-118) of the monthly subnetworks, 87 were associations among bacteria (Supplementary Table 2).

Although the temporal recurrence of associations over the ten years was low, we found high recurrence in corresponding months from different years. We quantified the fraction of subnetworks in which each association appeared (Supplementary Figure 6). We observed the highest prevalence from December to March, and the lowest prevalence from June to August (Supplementary Figure 6). For each month, we taxonomically characterized prevalent associations appearing in at least nine out of the ten monthly subnetworks (e.g., 9 out of 10 Januarys; Figure 3). We found a larger number of prevalent associations in colder waters compared to warmer waters, with Alphaproteobacteria dominating these associations, especially in April and May (Figure 3). The Alphaproteobacteria ASVs featuring highly prevalent associations belonged to *Pelagibacter ubique* (SAR11 Clades Ia & II), Rhodobacteraceae, *Amylibacter,* Puniceispirillales (SAR116), *Ascidiaceihabitans, Planktomarina,* Parvibaculales (OCS116), and *Kiloniella.* Between April and May, we noticed a large increase in the fraction of associations including Cyanobacteria or Bacteroidetes as association partners. While Cyanobacteria associations were a small fraction during November-April, they had a dominant role from May-October along with Bacteroidetes and Alphaproteobacteria associations (Figure 3). Overall, this underlines the dynamic nature of associations over the year, pointing to recurring annual associations that may be essential for ecosystem function.

**Figure 3:**
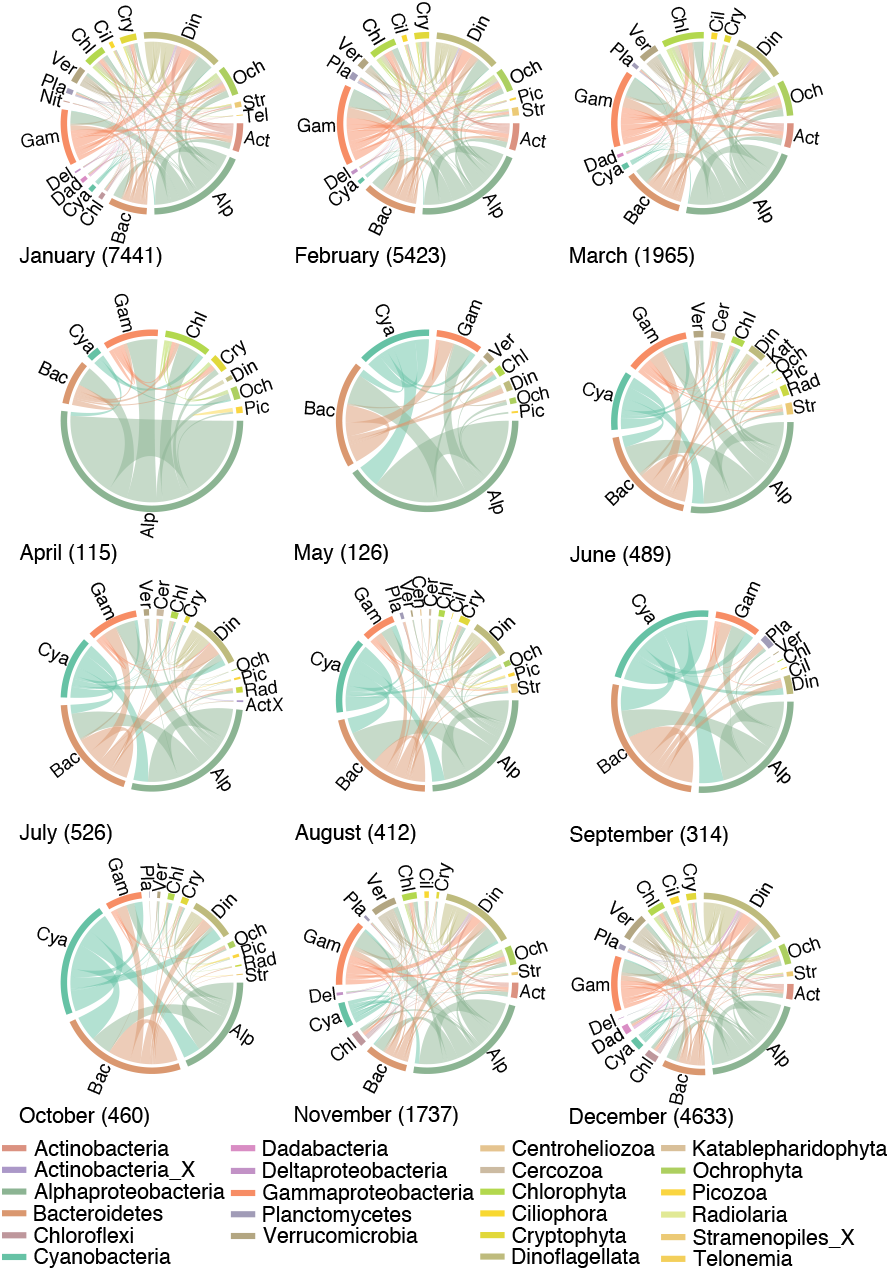
Associations with a monthly prevalence of at least 90%. Bacteria and eukaryotes are separated and ordered alphabetically. We provide in parentheses the number of associations that appeared in at least nine out of ten monthly subnetworks.

### Dynamic associations within main taxonomic groups: the case of Cyanobacteria

Our results indicated that associations are dynamic within specific taxonomic groups. Therefore, we investigated their behavior in Cyanobacteria given the importance of this group as primary producers in the ocean. We found 661 associations for *Synechococcus*, *Prochlorococcus,* and *Cyanobium* ASVs (Figure 4 and Supplementary Figure 7). Most associations between cyanobacterial ASVs were positive (63 of 65), and only a *Synechococcus* (referred to as bn_ASV_5) was negatively associated (association score −0.5) with other *Synechococcus* (bn_ASV_1 and bn_ASV_25), which, in turn, were positively associated (association score 0.8). While bn_ASV_5 appeared mainly in colder months, the other two appeared mainly in warmer months (Supplementary Figure 7). All Cyanobacteria had more associations with other bacteria (in total 433) than with eukaryotes (in total 163).

**Figure 4:**
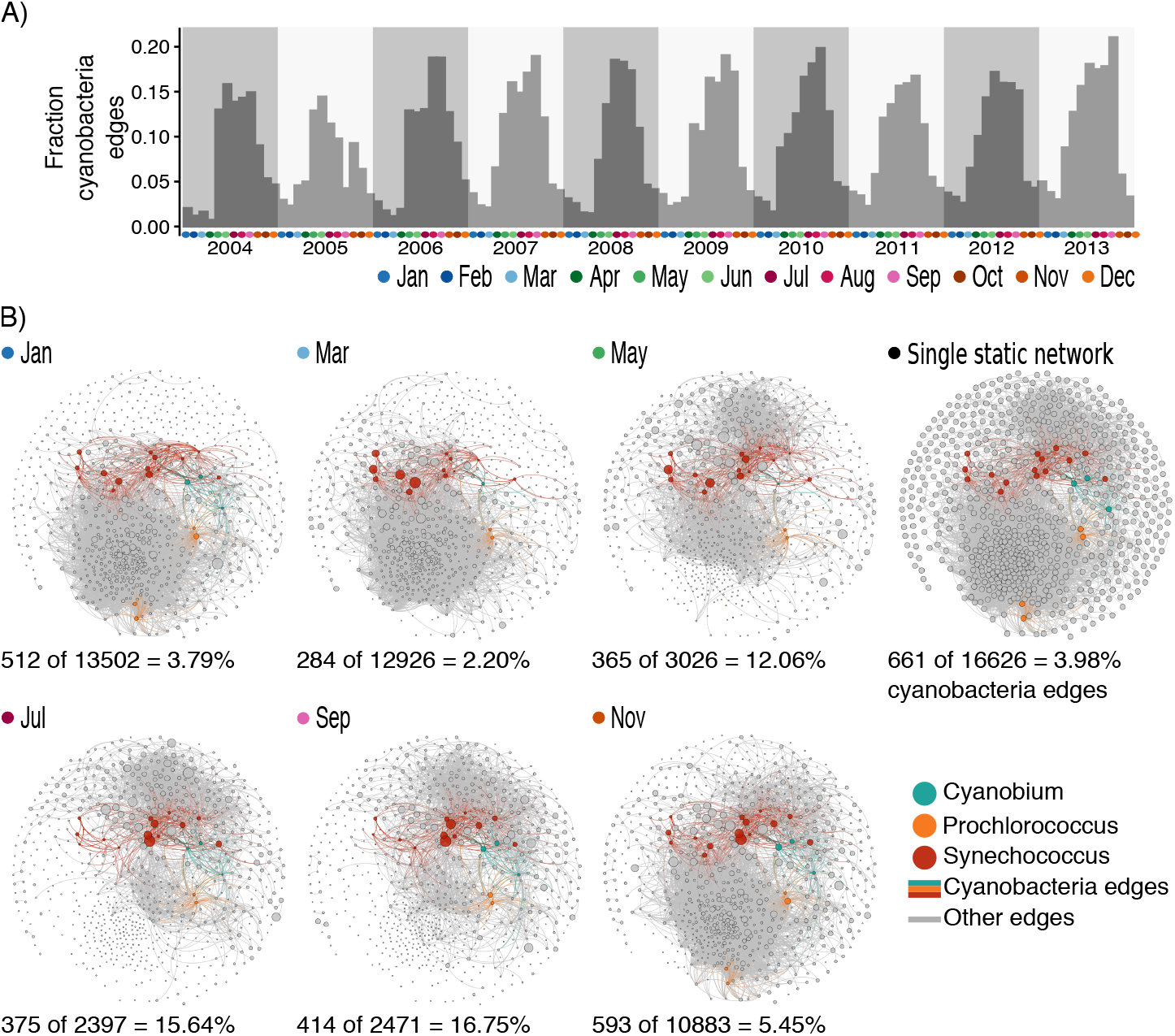
*Cyanobacteria* associations. A) Fraction of edges in the temporal network containing at least one *Cyanobacteria* ASV. B) Location of *Cyanobacteria* associations in the temporal network and the single static network. Here we show, as an example, selected months of year 2011. The number and fraction of cyanobacterial edges and total number of edges is listed below each monthly subnetwork and the single static network.

Within the temporal network, the fraction of Cyanobacteria associations was highest in April-October (Figure 4A), which are the months with the largest cyanobacterial abundances (Supplementary Figure 7), and the fewest edges in the entire temporal network (Figure 2B), for example, in the year 2011 (Figure 4B). We found that cyanobacterial ASVs, although being evolutionarily related, behaved differently in terms of the number of associations over time, and association partners (Supplementary Figure 7). For example,

*Synechococcus* bn_ASV_5 had fewer partners than bn_ASV_1 according to numbers of associations, but more according to taxonomic variety (Supplementary Figure 7). Only a tiny fraction of *Prochlorococcus* (e.g. bp_ASV_18) association partners were other Cyanobacteria, which contrasted with *Synechococcus* and *Cyanobium* (Supplementary Figure 7). Moreover, we observed that *Cyanobium* (bn_ASV_20) connected to one Deltaproteobacteria (SAR324) ASV during the first eight years, but the association disappeared in the last two years. In particular, the inferred association duration was 101 months, starting in March 2004 and ending in July 2012. After summer 2012, the Deltaproteobacteria ASV was not detected except a few reads in November and December 2012 and 2013. This Cyanobacteria example may also illustrate the dynamics of associations within other main taxonomic groups.

### Validating associations using known ecological interactions

We checked how many potential interactions could be validated using a database of observed ecological interactions (PIDA; [17]). In total, 16 associations (out of 16626) in the temporal network were validated by PIDA (Supplementary Table 6). These 16 associations describe six unique interactions between seven taxa (at the genus-level). For instance, the reoccurring association between a diatom from genus *Thalassiosira* and a Flavobacteriia starts mainly around October and often ends around March (Supplementary Figure 8). In contrast, the reoccurring association between a dinoflagellate from genus *Gyrodinium* and one from *Heterocapsa* appears for a shorter time and during the summer months (Supplementary Figure 8).

## DISCUSSION

Previous work identified yearly recurrence of microbial community composition at the BBMO [13, 28, 64], and similarly at the nearby Bay of Banyuls [14], both in the North-West Mediterranean Sea and in other temperate sites around the world [12, 65]. We focused here in the connectivity of microorganisms and how they organize themselves from a network perspective. In general, the measured global network metrics (edge density, transitivity, and average path length) are within the range reported in previous studies [22–25, 66–68] (Table 2). Contrary to early studies reporting biological networks generally being disassortative (negative assortativity based on degree) [69], our single static network and the monthly subnetworks were assortative. Microorganisms had more and stronger connections and a tighter clustering in colder than in warmer waters. To some extent, this might reflect species richness, which has been shown for the resident microorganisms to increase during the colder months at BBMO using the same dataset [28]. However, the exact effect of richness on ecological interactions among microorganisms needs further analysis. Seasonal bacterial freshwater networks [67] also showed higher clustering in fall and winter than in spring and summer, but, in contrast to our results, networks were most extensive in summer and smallest in winter. In agreement with our results, Chaffron et al. [68] reported higher association strength, edge density, and transitivity in cold polar regions compared to other warmer regions of the global ocean. Colder waters in the Mediterranean Sea are milder than polar waters. However, together, these results suggest that either microorganisms interact more in colder environments, or that their recurrence is higher due to higher environmental selection exerted by low temperatures. Additionally, limited resources (mainly nutrients) in summer or in the tropical and subtropical open ocean may prevent the establishment of several microbial interactions. In any case, temperature is likely not the only driver of network architecture [68].

**Table 2:** Global network metrics of previously described microbial association networks.

| Location & Depth | Edge density | Transitivity | Average path length | Sampling | Domains | Notes | Reference |
| --- | --- | --- | --- | --- | --- | --- | --- |
| SPOT (Off the southern California coast); Deep chlorophyll maximum. | 0.04 | 0.26 | 3.05 | Monthly. August 2000 - March 2004 | Archaea, bacteria, and eukaryotes | Edge density for microbial network including environmental factors. Transitivity and average path length for microbial network. | [22] |
| SPOT; Surface ocean and deep chlorophyll maximum | 0.14 | 0.33 | 1.94 | Monthly. August 2000 - January 2011 | Free-living bacteria and picoeukaryotes | Metrics from surface layer network. | [23] |
| SPOT; Surface | 0.02 | 0.24 |  | Monthly. March 2008 - January 2011 | Free-living eukaryotes (0.7–20 µm), bacteria (0.22–1 µm) and viruses (30 kDa–0.22 µm) |  | [24] |
| SPOT; Five depths (5 m - Surface, the deep chlorophyll maximum layer, 150 m, 500 m and 890 m - just above the sea floor) | 0.04 | 0.28 | 2.07 | Monthly. August 2003 - January 2011 | Free-living bacteria | Metrics for 5 m layer network. | [25] |
| 52 samples from freshwater lakes in China; Surface | (0.023)<br>W:0.033,<br>Sp:0.032<br>,<br>S:0.036,<br>F:0.029 | (0.472)<br>W:0.518,<br>Sp:0.480,<br>S:0.475,<br>F:0.573 | (4.84)<br>W:2.16,<br>Sp:5.03<br>S:7.26,<br>F:3.04 | Spatial | Bacteria | Metrics for (whole network) and seasonal networks: W: winter, Sp: spring, S: summer, and F: fall | [67] |
| 68 stations from the Tara Oceans expedition across eight oceanic provinces; Surface and Deep chlorophyll maximum | 0.005,<br>0.003,<br>0.008 | 0.2,<br>0.0,<br>0.43 | 3.05,<br>3.02,<br>2.56 | Spatial | Organisms from seven size fractions | Metrics from surface networks including eukaryotes only, eukaryotes and prokaryotes (0.5-5 µm), and prokaryotes only (0.2-1.6 µm) | [66] |
| 115 stations from the Tara Oceans expedition covering all major oceanic provinces from pole to pole; Surface and deep chlorophyll maximum | 0.002 | 0.036 |  | Spatial | Bacteria, archaea, and eukaryotes from six size fractions. | Metrics represent the means of sample-specific subnetworks. | [68] |

The effects of environmental variables on network metrics are unclear [70], yet, our approach allowed us to identify potential environmental drivers of network architecture. Correlation analyses pointed to variables that have been found to influence microbial abundances in the ocean. For instance, our results indicated that temperature and day length, key variables driving microbial assemblages in seasonal time series [12–14], and to a lesser extent inorganic nutrients, were the main factors influencing global network metrics. It also agrees with earlier works indicating that phosphorus and nitrogen are the primary limiting nutrients in the Western Mediterranean Sea [71, 72]. Altogether, our correlation analysis is a step forward towards elucidating the effects of environmental variables on network metrics. However, we did not consider several other variables that could affect network architecture (e.g. organic matter).

Our preliminary network (significant associations derived with eLSA) contained 18% negative edges compared to 0.9% in the single static network (after filtering). Thus, our filtering strategy removed proportionally more negative edges. Associations may represent positive or negative interactions, but they can also indicate high niche overlap (positive association) or divergent niches (negative association) between microorganisms [73]. We hypothesize that most of the removed negative edges represented associations between microorganisms from divergent niches, most likely corresponding to colder or warmer months.

We found more highly prevalent associations within specific months than when considering all ten years of data. Furthermore, our results indicate a potentially low number of core interactions and a vast number of non-core ones. Usually, core microorganisms are defined based on sequence abundances, as those microorganisms (or taxonomical groups) appearing in all samples or habitats being under investigation [74]. Shade & Handelsman [74] suggested that other parameters, including connectivity, should create a more complex portrait of the core microbiome and advance our understanding of the role of key microorganisms and functions within and across ecosystems [74]. Using a temporal network we identified core associations based on recurrence, which contributes to our understanding of key interactions underpinning microbial ecosystem functions. Considering associations within each month, we found more highly-prevalent associations in colder than in warmer months. Our results indicate microbial connectivity is more repeatable (indicating higher predictability) in colder than in warmer waters. On the one hand, the microbial community in colder waters being more recurrent [13] may explain our observations indicating a more robust connectivity during this period. Alternatively, it may be the stronger connectivity what leads to more similar communities in colder waters at the BBMO. Last but not least, the interplay of both species dynamics and interactions may determine community turnover in the studied ecosystem. From a technical viewpoint, our monthly sampling strategy and/or the overall single static network may have not been able to detect interactions appearing solely in summer resulting in smaller monthly subnetworks. For instance, previous work on freshwater lakes constructed season-specific networks and found more associations in summer than in winter [67].

Several network-based analyses have been used to particularly study Cyanobacteria associations. For example, in the southern Californian coast, Chow et al. [24] identified 44 potential relationships of 12 Cyanobacteria (*Prochlorococcus* and *Synechococcus*) with two potential eukaryote grazers (a ciliate and a dinoflagellate), 39 to other bacteria, and three between Cyanobacteria, which were all positive. Similarly, all cyanobacterial ASVs in our study connected primarily to other bacterial ASVs, and featured mainly positive associations. Furthermore, Cyanobacteria displayed primarily positive associations with other microorganisms in a global ocean network [66]. This suggests that other sampling or computational approaches are needed to detect negative associations involving marine cyanobacteria.

Identifying different potential association partners for closely related Cyanobacteria may indicate adaptations to different niches. A recent study found distinct seasonal patterns for closely related bacterial taxa indicating niche partitioning at the BBMO, including *Synechococcus* ASVs [64]. Our approach can complement and further characterize “subniches” by providing association partners for different ASVs. Moreover, in contrast to a single static network, temporal networks allow identifying associated partners in time (Supplementary Figure 7). An increase in the abundance of a microorganism may promote the growth of associated partners and a decrease may hinder the growth of partners or cause predators to prey on other microorganisms. Moreover, given the majority of association partners being other bacteria, the growth of Cyanobacteria may affect other bacteria and their growth, which is why it is necessary to identify potential interaction partners [67].

Our approach allowed us to disentangle in time the associations captured by a single static network using monthly samples for ten years. Future studies should determine whether higher sampling frequency (e.g., daily samples during a month) can capture other associations not present in our networks. Thus, our results should be considered taking into account the used (monthly) sampling frequency. In addition, certain network metrics may depend on the tool used to infer the single static network, e.g., edge density, and, therefore, should be interpreted with care. An additional consideration is that we disregarded local network patterns by using global network metrics. Future work could use the local-topological metric based on graphlets [75]. Counting the number of graphlets a node is part of quantifies their local connection patterns, which allows inferring seasonal microorganisms through recurring connection patterns in a temporal network.

## CONCLUSION

Incorporating the temporal dimension in microbial association analysis unveiled multiple patterns that often remain hidden when using single static networks. Investigating a coastal marine microbial ecosystem over ten years revealed a one-year-periodicity in the network topology. The temporal network architecture was not stochastic, but displayed a modest amount of recurrence over time, especially in winter. Future efforts to understand the ocean microbiome should consider the dynamics of microbial interactions as these are likely fundamental for ecosystem functioning.

## Ethics approval and consent to participate

Not applicable.

## Consent for publication

Not applicable.

## Availability of data and material

The BBMO microbial sequence abundances (ASV tables), taxonomic classifications, environmental data including nutrients, networks and R-Markdowns for data analysis including commands to run eLSA and EnDED (environmentally-driven-edge-detection and computing Jaccard index) are publicly available: https://github.com/InaMariaDeutschmann/TemporalNetworkBBMO.

## Author’s contributions

The overall project was conceived and designed by RL and AKK. VB, JMG, and RM were responsible for the sampling and contextual data at the BBMO. RL processed the amplicon data from BBMO generating the ASV tables. AKK constructed the initial preliminary network which was the starting point of the present study. IMD developed the conceptual approach and DE, SC, and RL contributed to its finalization. IMD performed the data analysis. ED, DE, SC, FL, AKK, CM, JMG, and RL contributed with the biological interpretation of the results. IMD wrote the original draft. All authors contributed to manuscript revisions and approved the final version of the manuscript.

## Acknowledgements

We thank all members of the Blanes Bay Microbial Observatory (http://bbmo.icm.csic.es) team with the multiple projects funding this collaborative effort. Part of the analyses have been performed at the Marbits bioinformatics core at ICM-CSIC (https://marbits.icm.csic.es). We thank L. Felipe Benites for fruitful discussions regarding microorganisms and their interactions.

## References

1. Falkowski PG, Fenchel T, Delong EF. The Microbial Engines That Drive Earth’s Biogeochemical Cycles. Science. 2008;320:1034–9.

2. DeLong EF. The microbial ocean from genomes to biomes. Nature. 2009;459:200–6.

3. Krabberød AK, Bjorbækmo MFM, Shalchian-Tabrizi K, Logares R. Exploring the oceanic microeukaryotic interactome with metaomics approaches. Aquatic Microbial Ecology. 2017;79:1–12.

4. Locey KJ, Lennon JT. Scaling laws predict global microbial diversity. Proceedings of the National Academy of Sciences. 2016;113:5970–5.

5. Kallmeyer J, Pockalny R, Adhikari RR, Smith DC, D’Hondt S. Global distribution of microbial abundance and biomass in subseafloor sediment. Proceedings of the National Academy of Sciences. 2012;109:16213–6.

6. Whitman WB, Coleman DC, Wiebe WJ. Prokaryotes: The unseen majority. Proceedings of the National Academy of Sciences. 1998;95:6578–83.

7. Bar-On YM, Milo R. The Biomass Composition of the Oceans: A Blueprint of Our Blue Planet. Cell. 2019;179:1451–4.

8. Bar-On YM, Phillips R, Milo R. The biomass distribution on Earth. Proceedings of the National Academy of Sciences. 2018;115:6506–11.

9. Vellend M. The theory of ecological communities (MPB-57). Princeton University Press; 2020.

10. Lindström ES, Langenheder S. Local and regional factors influencing bacterial community assembly. Environmental Microbiology Reports. 2012;4:1–9.

11. Mori AS, Isbell F, Seidl R. β-Diversity, Community Assembly, and Ecosystem Functioning. Trends in Ecology & Evolution. 2018;33:549–64.

12. Bunse C, Pinhassi J. Marine Bacterioplankton Seasonal Succession Dynamics. Trends in Microbiology. 2017;25:494–505.

13. Giner CR, Balagué V, Krabberød AK, Ferrera I, Reñé A, Garcés E, et al. Quantifying long-term recurrence in planktonic microbial eukaryotes. Molecular Ecology. 2019;28:923–35.

14. Lambert S, Tragin M, Lozano J-C, Ghiglione J-F, Vaulot D, Bouget F-Y, et al. Rhythmicity of coastal marine picoeukaryotes, bacteria and archaea despite irregular environmental perturbations. The ISME Journal. 2019;13:388.

15. Logares R, Deutschmann IM, Junger PC, Giner CR, Krabberød AK, Schmidt TSB, et al. Disentangling the mechanisms shaping the surface ocean microbiota. Microbiome. 2020;8:55.

16. Barraclough TG. How Do Species Interactions Affect Evolutionary Dynamics Across Whole Communities? Annu Rev Ecol Evol Syst. 2015;46:25–48.

17. Bjorbækmo MFM, Evenstad A, Røsæg LL, Krabberød AK, Logares R. The planktonic protist interactome: where do we stand after a century of research? The ISME Journal. 2019. https://doi.org/10.1038/s41396-019-0542-5.

18. Baldauf SL. An overview of the phylogeny and diversity of eukaryotes. Journal of Systematics and Evolution. 2008;46:263.

19. Lewis WH, Tahon G, Geesink P, Sousa DZ, Ettema TJG. Innovations to culturing the uncultured microbial majority. Nature Reviews Microbiology. 2020. https://doi.org/10.1038/s41579-020-00458-8.

20. Weiss S, Van Treuren W, Lozupone C, Faust K, Friedman J, Deng Y, et al. Correlation detection strategies in microbial data sets vary widely in sensitivity and precision. The ISME Journal. 2016;10:1669–81.

21. Layeghifard M, Hwang DM, Guttman DS. Disentangling Interactions in the Microbiome: A Network Perspective. Trends in Microbiology. 2017;25:217–28.

22. Steele JA, Countway PD, Xia L, Vigil PD, Beman JM, Kim DY, et al. Marine bacterial, archaeal and protistan association networks reveal ecological linkages. The ISME Journal. 2011;5:1414–25.

23. Chow C-ET, Sachdeva R, Cram JA, Steele JA, Needham DM, Patel A, et al. Temporal variability and coherence of euphotic zone bacterial communities over a decade in the Southern California Bight. The ISME Journal. 2013;7:2259–73.

24. Chow C-ET, Kim DY, Sachdeva R, Caron DA, Fuhrman JA. Top-down controls on bacterial community structure: microbial network analysis of bacteria, T4-like viruses and protists. The ISME Journal. 2014;8:816–29.

25. Cram JA, Xia LC, Needham DM, Sachdeva R, Sun F, Fuhrman JA. Cross-depth analysis of marine bacterial networks suggests downward propagation of temporal changes. The ISME Journal. 2015;9:2573–86.

26. Needham DM, Sachdeva R, Fuhrman JA. Ecological dynamics and co-occurrence among marine phytoplankton, bacteria and myoviruses shows microdiversity matters. The ISME Journal. 2017;11:1614–29.

27. Parada AE, Fuhrman JA. Marine archaeal dynamics and interactions with the microbial community over 5 years from surface to seafloor. The ISME Journal. 2017;11:2510–25.

28. Krabberød AK, Deutschmann IM, Bjorbækmo MFM, Balagué V, Giner CR, Ferrera I, et al. Long-term patterns of an interconnected core marine microbiota. Environmental Microbiome. 2022;17:22.

29. Blonder B, Wey TW, Dornhaus A, James R, Sih A. Temporal dynamics and network analysis. Methods in Ecology and Evolution. 2012;3:958–72.

30. Gasol JM, Cardelús C, G Morán XA, Balagué V, Forn I, Marrasé C, et al. Seasonal patterns in phytoplankton photosynthetic parameters and primary production at a coastal NW Mediterranean site. Scientia Marina. 2016;80:63–77.

31. Ferrera I, Reñé A, Funosas D, Camp J, Massana R, Gasol JM, et al. Assessment of microbial plankton diversity as an ecological indicator in the NW Mediterranean coast. Marine Pollution Bulletin. 2020;160:111691.

32. Grasshoff K, Kremling K, Ehrhardt M. Methods of seawater analysis. John Wiley & Sons; 2009.

33. Schauer M, Balagué V, Pedrós-Alió C, Massana R. Seasonal changes in the taxonomic composition of bacterioplankton in a coastal oligotrophic system. Aquatic Microbial Ecology. 2003;31:163–74.

34. Stoeck T, Bass D, Nebel M, Christen R, Jones MDM, Breiner H-W, et al. Multiple marker parallel tag environmental DNA sequencing reveals a highly complex eukaryotic community in marine anoxic water. Molecular Ecology. 2010;19:21–31.

35. Herlemann DP, Labrenz M, Jürgens K, Bertilsson S, Waniek JJ, Andersson AF. Transitions in bacterial communities along the 2000 km salinity gradient of the Baltic Sea. The ISME Journal. 2011;5:1571–9.

36. Apprill A, McNally S, Parsons R, Weber L. Minor revision to V4 region SSU rRNA 806R gene primer greatly increases detection of SAR11 bacterioplankton. Aquatic Microbial Ecology. 2015;75:129–37.

37. Callahan BJ, McMurdie PJ, Rosen MJ, Han AW, Johnson AJA, Holmes SP. DADA2: High-resolution sample inference from Illumina amplicon data. Nature Methods. 2016;13:581–3.

38. Oksanen J, Blanchet FG, Friendly M, Kindt R, Legendre P, McGlinn D, et al. vegan: Community Ecology Package. 2019.

39. Moritz S, Gatscha S. imputeTS: Time Series Missing Value Imputation. 2017.

40. Wang Q, Garrity GM, Tiedje JM, Cole JR. Naïve Bayesian Classifier for Rapid Assignment of rRNA Sequences into the New Bacterial Taxonomy. Applied and Environmental Microbiology. 2007;73:5261–7.

41. Quast C, Pruesse E, Yilmaz P, Gerken J, Schweer T, Yarza P, et al. The SILVA ribosomal RNA gene database project: improved data processing and web-based tools. Nucleic Acids Research. 2012;41:D590–6.

42. Altschul SF, Gish W, Miller W, Myers EW, Lipman DJ. Basic local alignment search tool. Journal of Molecular Biology. 1990;215:403–10.

43. Guillou L, Bachar D, Audic S, Bass D, Berney C, Bittner L, et al. The Protist Ribosomal Reference database (PR$^2$): a catalog of unicellular eukaryote Small Sub-Unit rRNA sequences with curated taxonomy. Nucleic Acids Research. 2012;41:D597–604.

44. McNichol J, Berube PM, Biller SJ, Fuhrman JA, Gilbert JA. Evaluating and Improving Small Subunit rRNA PCR Primer Coverage for Bacteria, Archaea, and Eukaryotes Using Metagenomes from Global Ocean Surveys. mSystems. 2021;6:e00565–21.

45. Balzano S, Abs E, Leterme SC. Protist diversity along a salinity gradient in a coastal lagoon. Aquat Microb Ecol. 2015;74:263–77.

46. Xia LC, Steele JA, Cram JA, Cardon ZG, Simmons SL, Vallino JJ, et al. Extended local similarity analysis (eLSA) of microbial community and other time series data with replicates. BMC Systems Biology. 2011;5:S15.

47. Xia LC, Ai D, Cram J, Fuhrman JA, Sun F. Efficient statistical significance approximation for local similarity analysis of high-throughput time series data. Bioinformatics. 2013;29:230–7.

48. Deutschmann IM, Lima-Mendez G, Krabberød AK, Raes J, Vallina SM, Faust K, et al. Disentangling environmental effects in microbial association networks. Microbiome. 2021;9:232.

49. R Core Team. R: A Language and Environment for Statistical Computing. Vienna, Austria: R Foundation for Statistical Computing; 2019.

50. Connor N, Barberán A, Clauset A. Using null models to infer microbial co-occurrence networks. PLOS ONE. 2017;12:1–23.

51. Poisot T, Canard E, Mouillot D, Mouquet N, Gravel D. The dissimilarity of species interaction networks. Ecology Letters. 2012;15:1353–61.

52. Csardi G, Nepusz T. The igraph software package for complex network research. InterJournal. 2006;Complex Systems:1695.

53. Jamakovic A, Uhlig S. On the relationships between topological measures in real-world networks. Networks & Heterogeneous Media. 2008;3:345–59.

54. Gu Z, Gu L, Eils R, Schlesner M, Brors B. circlize implements and enhances circular visualization in R. Bioinformatics. 2014;30:2811–2.

55. Holm S. A Simple Sequentially Rejective Multiple Test Procedure. Scandinavian Journal of Statistics. 1979;6:65–70.

56. Revelle W. psych: Procedures for Psychological, Psychometric, and Personality Research. Evanston, Illinois: Northwestern University; 2020.

57. Bastian M, Heymann S, Jacomy M. Gephi: An Open Source Software for Exploring and Manipulating Networks. ICWSM. 2009;3.

58. Fruchterman TMJ, Reingold EM. Graph drawing by force-directed placement. Software: Practice and Experience. 1991;21:1129–64.

59. Gloor GB, Macklaim JM, Pawlowsky-Glahn V, Egozcue JJ. Microbiome Datasets Are Compositional: And This Is Not Optional. Frontiers in Microbiology. 2017;8:2224.

60. Hirano H, Takemoto K. Difficulty in inferring microbial community structure based on co-occurrence network approaches. BMC Bioinformatics. 2019;20:329.

61. Tackmann J, Rodrigues JFM, von Mering C. Rapid Inference of Direct Interactions in Large-Scale Ecological Networks from Heterogeneous Microbial Sequencing Data. Cell Systems. 2019;9:286–296.e8.

62. Garczarek L, Guyet U, Doré H, Farrant GK, Hoebeke M, Brillet-Guéguen L, et al. Cyanorak v2.1: a scalable information system dedicated to the visualization and expert curation of marine and brackish picocyanobacteria genomes. Nucleic Acids Research. 2021;49:D667–76.

63. Mestre M, Höfer J, Sala MM, Gasol JM. Seasonal Variation of Bacterial Diversity Along the Marine Particulate Matter Continuum. Frontiers in Microbiology. 2020;11:1590.

64. Auladell A, Barberán A, Logares R, Garcés E, Gasol JM, Ferrera I. Seasonal niche differentiation among closely related marine bacteria. The ISME Journal. 2022;16:178–89.

65. Fuhrman JA, Cram JA, Needham DM. Marine microbial community dynamics and their ecological interpretation. Nature Reviews Microbiology. 2015;13:133–46.

66. Lima-Mendez G, Faust K, Henry N, Decelle J, Colin S, Carcillo F, et al. Determinants of community structure in the global plankton interactome. Science. 2015;348:1262073.

67. Zhao D, Shen F, Zeng J, Huang R, Yu Z, Wu QL. Network analysis reveals seasonal variation of co-occurrence correlations between Cyanobacteria and other bacterioplankton. Science of The Total Environment. 2016;573:817–25.

68. Chaffron S, Delage E, Budinich M, Vintache D, Henry N, Nef C, et al. Environmental vulnerability of the global ocean epipelagic plankton community interactome. Sci Adv. 2021;7.

69. Newman MEJ. Assortative Mixing in Networks. Phys Rev Lett. 2002;89:208701.

70. Röttjers L, Faust K. From hairballs to hypotheses–biological insights from microbial networks. FEMS Microbiology Reviews. 2018;42:761–80.

71. Estrada M. Primary production in the northwestern Mediterranean. 1996.

72. Sala MM, Peters Francesc, Gasol JM, Pedrós-Alió C, Marrasé C, Vaqué D. Seasonal and spatial variations in the nutrient limitation of bacterioplankton growth in the northwestern Mediterranean. Aquatic Microbial Ecology. 2002;27:47–56.

73. Hernandez DJ, David AS, Menges ES, Searcy CA, Afkhami ME. Environmental stress destabilizes microbial networks. The ISME Journal. 2021. https://doi.org/10.1038/s41396-020-00882-x.

74. Shade A, Handelsman J. Beyond the Venn diagram: the hunt for a core microbiome. Environmental Microbiology. 2012;14:4–12.

75. Pržulj N, Corneil DG, Jurisica I. Modeling interactome: scale-free or geometric? Bioinformatics. 2004;20:3508–15.

