## Supplementary Material for "Disentangling temporal associations in marine microbial networks"

#### FIGURES LEGENDS

**Supplementary Figure 1:** Correlation Analysis. Using the temporal network, we correlated six global network metrics with environmental factors including the nutrients $PO_4^{3-}$, $NH_4^{+}$, $NO_2^{-}$, $NO_3^{-}$, and $SiO_2$. The global network metrics were: Edge density, Average positive association (Avg. pos. ass.) score, Transitivity, Average path length (Avg. path length), Assortativity (degree), and Assortativity (bacteria vs. eukaryote). Each dot is a sample-specific subnetwork and its color indicates the month it represents. Also, the linear regression line with a 0.95 confidence interval is shown in grey.

**Supplementary Figure 2:** Correlation Analysis through linear regression. Using the temporal network, we correlated six global network metrics with environmental factors including the nutrients $PO_4^{3-}$, $NH_4^{+}$, $NO_2^{-}$, $NO_3^{-}$, and $SiO_2$. The global network metrics were: Edge density, Average positive association (Avg. pos. ass.) score, Transitivity, Average path length (Avg. path length), Assortativity (degree), and Assortativity (bacteria vs. eukaryote). The number, circle's size, and color in the square correspond to the Spearman correlation scores, no circle indicates non-significance.

**Supplementary Figure 3:** Global (sub)network metrics. Number of nodes, number of edges, and six selected global network metrics for each sample-specific subnetwork of the temporal network determined with FlashWeave.

**Supplementary Figure 4:** Number of preserved, gained, and lost edges in summer and winter. A) Indicates how we determined summer indicated with red dots (temperature above 17 ºC and day length above 14 hours) and winter indicated with blue dots (temperature below 17 ºC and day length below 11 hours); grey dots indicate months that are neither summer nor winter. B) accumulation curve of ASVs per year for winter (blue) and summer (red). C) and D) the number of preserved, gained, and lost edges for winter and summer, respectively. The colors of flows indicate the prevalence of an edge with 10 (light blue) being present in each year, and 1 (dark blue) appearing in only one year. An edge appears in a year if it appears in at least one monthly subnetwork in the corresponding season. In winter, most edges appear in all years (light blue indicating 100% prevalence with edges present in all ten years), i.e. most edges are preserved in the consecutive months (we see a flow from the blue preserved box to the next blue box). In summer, compared to winter, fewer edges are present in a month (combination of boxes indicating preserved, first time gained, and gained), and more edges are (re)gained and lost throughout the years (subsequently prevalence is lower indicated through darker blue).

**Supplementary Figure 5:** Association prevalence increases slightly when microorganisms are taxonomically more related. We grouped the associations according to the taxonomic classification of association partners (columns) and size fractions (rows). For example "Class" groups associations between bacteria and eukaryotes, respectively, which were assigned to the same class. The grey column groups associations between bacteria and eukaryotes. The boxplot shows the association prevalence over a decade, i.e. in how many monthly subnetworks an association appears (given as a fraction from 0 to 100% = 120 networks).

**Supplementary Figure 6:** Association prevalence per month. Big bar plots: distribution of associations' prevalence for each month. For example, the bar at 100 for January indicates the number of edges that have been present in all Januarys of the ten-year time series. Small bar plots: number of nodes forming the associations with a 100% prevalence. For example, only bacteria were responsible for the edges during May, with an association prevalence of 100%. Bacteria are indicated with B or b, eukaryote with E or e. ASVs from the nano size-fraction have a capital letter (B, E), and ASVs from the pico size-fraction have a small letter (b, e).

**Supplementary Figure 7:** Association Partners of Cyanobacteria. The number of Cyanobacteria associations in the temporal network (stacked bars) and the cyanobacterial sequence abundance in each month (black dashed line). Within the box, figures are split by ASVs (rows) and size fractions: picoplankton (left column) and nanoplankton (right column). The unboxed plots on the right are ASVs detected only in the nanoplankton. The height of the bar indicates the number of edges in each month for each cyanobacterial ASV. The color indicates the taxonomy of the association partner. From bottom to top, first appear bacteria, and then eukaryotes, both sorted alphabetically. The subtitle shows the number of association partners followed by an identifier (first 3 letters) for bacteria and eukaryotes.

**Supplementary Figure 8:** Microbial association partners that have been reported in the literature. Found associations in the temporal network (one association per panel and a black dot on the bottom shows presence in the monthly subnetwork) and the sequence abundance in each month (solid and dashed lines). The color and line type indicate the taxonomy of the association partners.

### SUPPLEMENTARY TABLES

**Supplementary Table 1**: Number of nodes, removed isolated nodes, and number and fraction of edges in the preliminary network (A), and network obtained after removing environmentally-driven edges (B) and edges with association partners appearing more often alone than with the partner (C), which is the single static network. For comparison, we also give the minimum and maximum number of nodes and edges for the temporal network (D). We did not determine the union and intersection for the temporal network. If an ASV appeared in the nano and pico size fraction, it is counted twice. Therefore, for A-C) we also determined the number of microorganisms not considering size fraction (union) and being present in both size fractions (both, i.e. intersection).

| | A) eLSA | B) EnDED | C) Static network | D) Range in Temporal network |
|---|---|---|---|---|
| Connected nodes | 754 | 754 | 709 | 130-542 |
| Bacteria (pico) | 169 | 169 | 164 | 13-148 |
| Bacteria (nano) | 279 | 279 | 251 | 31-204 |
| Bacteria (union) | 309 | 309 | 281 | |
| Bacteria (both) | 139 | 139 | 134 | |
| Eukaryote (pico) | 150 | 150 | 141 | 7-124 |
| Eukaryote (nano) | 156 | 156 | 153 | 2-138 |
| Eukaryote (union) | 306 | 306 | 294 | |
| Eukaryote (both) | 0 | 0 | 0 | |
| Isolated nodes | 1000 | 0 | 45 | 6-38 |
| Edges | 29820 | 26505 | 16626 | 538-15083 |
| Positive edges | 24458 | 23405 | 16481 | 523-14940 |
| (%) | 82.0 | 88.3 | 99.1 | 92.2-99.7 |
| Negative edges | 5362 | 3100 | 145 | 12-143 |
| (%) | 18.0 | 11.7 | 0.9 | 0.3-7.8 |

*pico and nano – microorganism detected in the picoplankton and nanoplankton, respectively, union – how many microorganisms when not considering size-fraction, both – how many microorganims appear in both size fractions*

**Supplementary Table 2**: Top 100 most prevalent/recurring associations

| Association partners | Number of associations |
|---|---|
| Bacterial association in picoplankton | 42 |
| Bacterial association in nanoplankton | 35 |
| Bacterial associations between size fractions | 10 |
| Bacteria associated to Eukaryote in nanoplankton | 4 |
| Eukaryotic association in nanoplankton | 3 |
| Bacteria associated to Eukaryote in picoplankton | 3 |
| Bacteria in nanoplankton associated to Eukaryotic picoplankton | 2 |
| Eukaryotic association in picoplankton | 1 |

**Supplementary Table 3**: Number of environmental factors leading to the removal of edges.

| Number of environmental factors | Edges | Positive edges | Negative edges |
|---|---|---|---|
| 0, i.e. not environmentally-driven edges | 26505 | 23405 (88.3%) | 3100 (11.7%) |
| 1 | 2747 | 1019 (37.1%) | 1728 (62.9%) |
| 2 | 506 | 33 (6.5%) | 473 (93.5%) |
| 3 | 61 | 1 (1.6%) | 60 (98.4%) |
| 4 | 1 | 0 (0%) | 1 (100%) |

**Supplementary Table 4**: Number of environmentally-driven edges for each environmental factor and fraction considering the total number of edges (29820) in the network. In addition, we present the number of positive and negative edges and the fraction considering the number of edges removed through an environmental factor.

| Environmental factor | Edges | Positive edges | Negative edges |
|---|---|---|---|
| Temperature | 1920 (6.44%) | 725 (37.8%) | 1195 (62.2%) |
| Total chlorophyll-a concentration | 838 (2.81%) | 82 (9.8%) | 756 (90.2%) |
| Day length | 730 (2.45%) | 237 (32.5%) | 493 (67.5%) |
| $NO_2^-$ | 192 (0.64%) | 26 (13.5%) | 166 (86.5%) |
| $SiO_2$ | 162 (0.54%) | 6 (3.7%) | 156 (96.3%) |
| $NO_3^-$ | 57 (0.19%) | 12 (21.1%) | 45 (78.9%) |
| Turbidity | 47 (0.16%) | 0 | 47 (100%) |
| Salinity, $NH_4^+$, and $PO_4^{3-}$ | 0 | 0 | 0 |

**Supplementary Table 5**: 100% Matching sequences from Cyanorak database for selected cyanobacterial ASVs

| ASV | Number | Matching sequence name with clade and subclade |
|---|---|---|
| Synechococcus #1 | 38 | 2x A15-24 III IIIa, 2x A15-28 III IIIb, 3x A15-44 II IIa, 2x A15-62 II IIc, 2x A18-40 III IIIa, 2x A18-46.1 III IIIa, 2x BOUM118 III IIIa, 2x CC9605 II IIc, 2x M16.1 II IIa, 2x PROS-U-1 II IIh, 2x ROS8604 I Ib, 3x RS9902 II IIa, 3x RS9907 II IIa, 2x RS9915 III IIIa, 2x TAK9802 II IIa, 1x WH8016 I Ib, 2x WH8103 III IIIa, 2x WH8109 II IIa |
| Synechococcus #5 | 2 | 2x PROS-9-1 I Ib |
| Prochlorococcus #18 | 2 | 1x EQPAC1 HLI HLI, 1x MED4 HLI HLI |
| Cyanobium #20 | 2 | 1x MINOS11 5.3 5.3, 1x RCC307 5.3 5.3 |

**Supplementary Table 6**: Interactions found in the BBMO temporal network that have been reported in the literature. The table shows the number of associations found in the network. For example, the association between the ASVs classified as *Dia. Thalassiosira* and ASVs classified as F. unknown *Flavobacteriia* has been found 6 times in the network.

| Microorganisms | Occurrences | ID in PIDA |
|---|---|---|
| Dia. *Thalassiosira* - F. unknown Flavobacteriia | 6 | 2199 |
| Dino. *Heterocapsa* - Dino. *Prorocentrum* | 1 | 1501, 1511 |
| Dino. *Gyrodinium* - Dino. *Heterocapsa* | 1 | 1313, 1314, 1780, 1783 |
| Dino. *Prorocentrum* - Dino. *Gymnodinium* | 2 | 1499 |
| Dino. *Prorocentrum* - Dino. *Prorocentrum* | 4 | 1509, 1510 |
| Dino. *Prorocentrum* - Dino. *Scrippsiella* | 2 | 1513 |

Abbreviations indicate Dia - Diatomea; Dino - Dinoflagellata; F - Flavobacteriia; ID in PIDA refers to the number PIDA gave to an interaction described in the literature.
