## Supplementary figures and images for "Disentangling temporal associations in marine microbial networks"

### Supplementary Figure 1

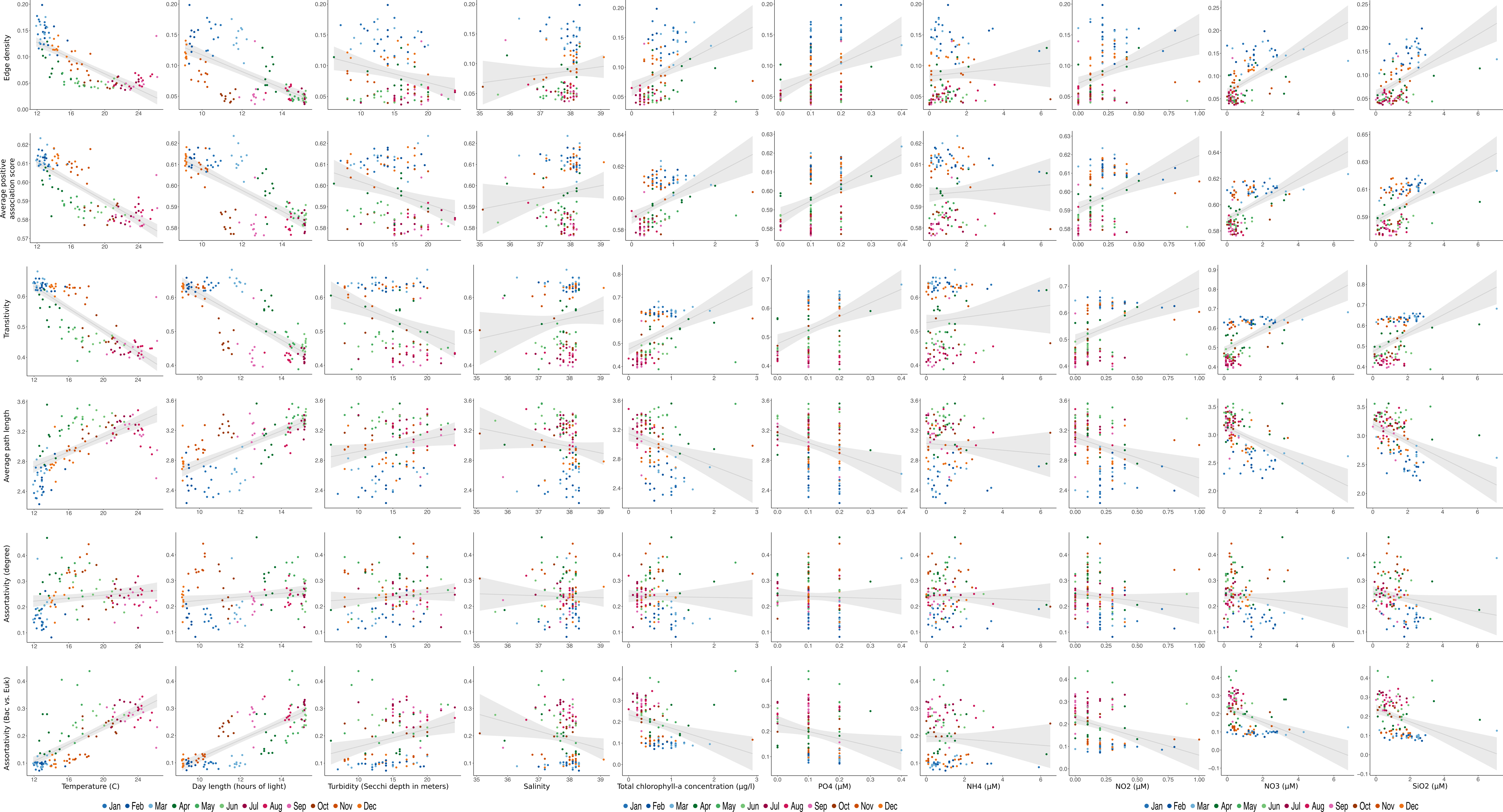

### Supplementary Figure 2

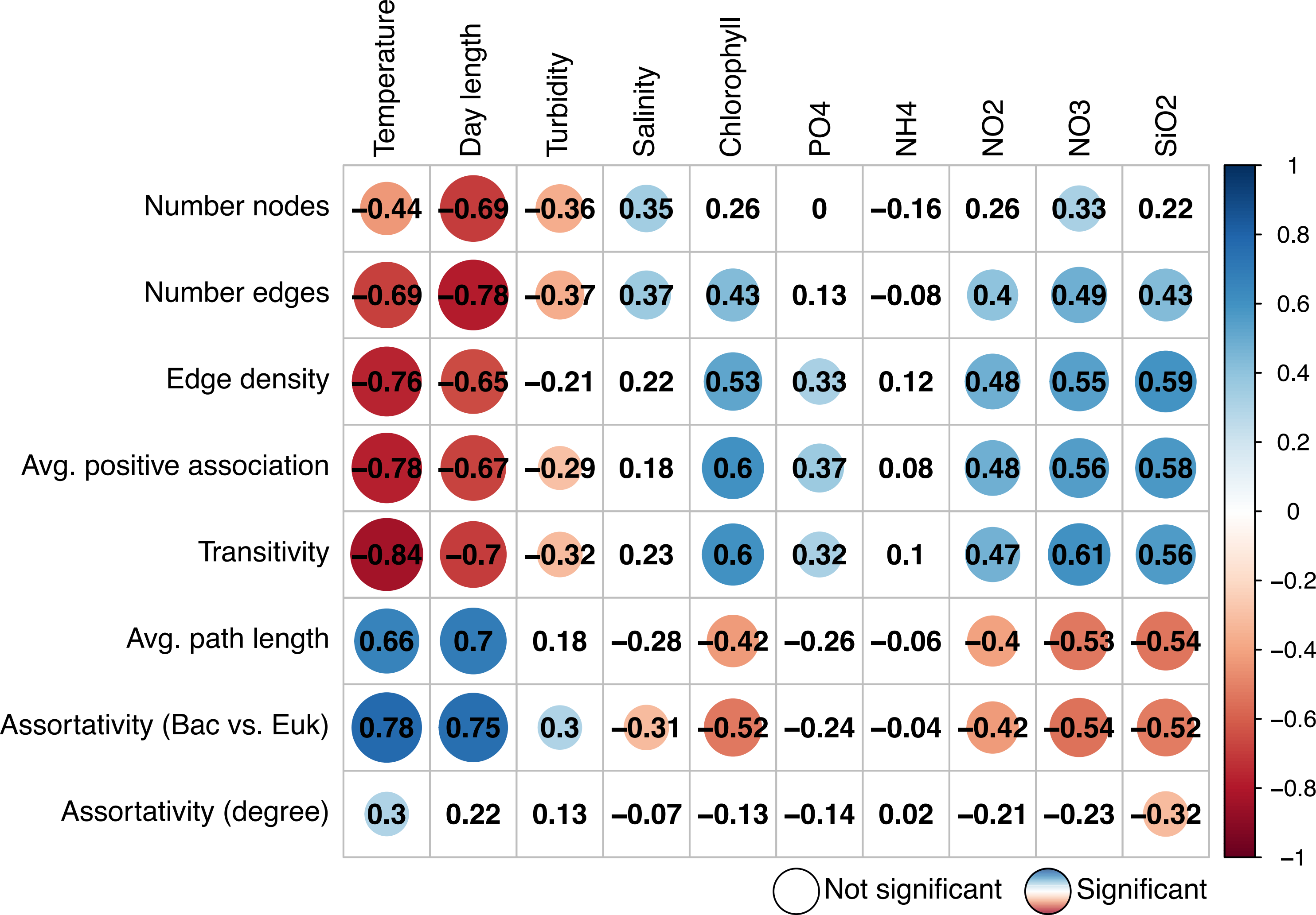

### Supplementary Figure 3

#### FlashWeave with environmental factors

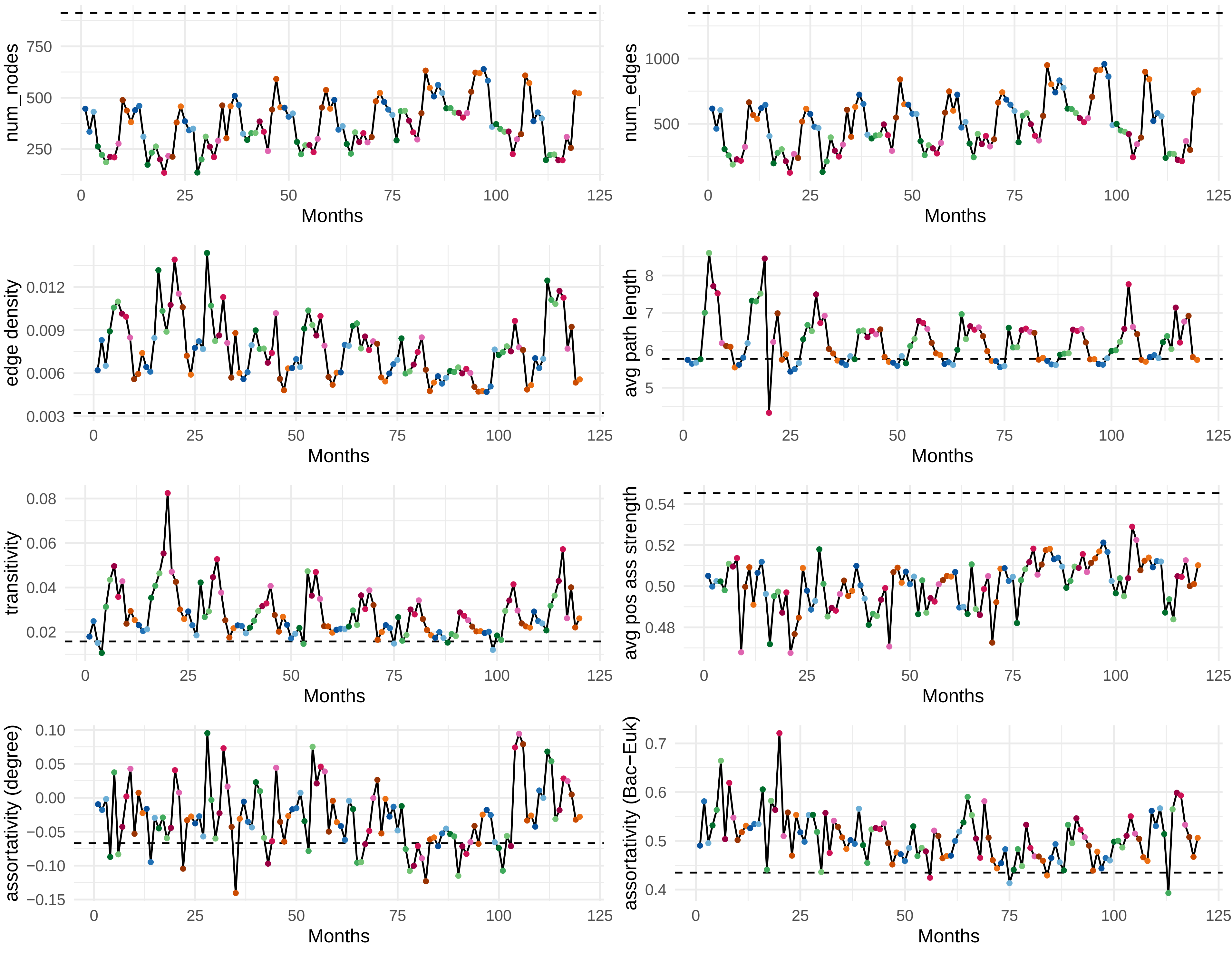

### Supplementary Figure 4

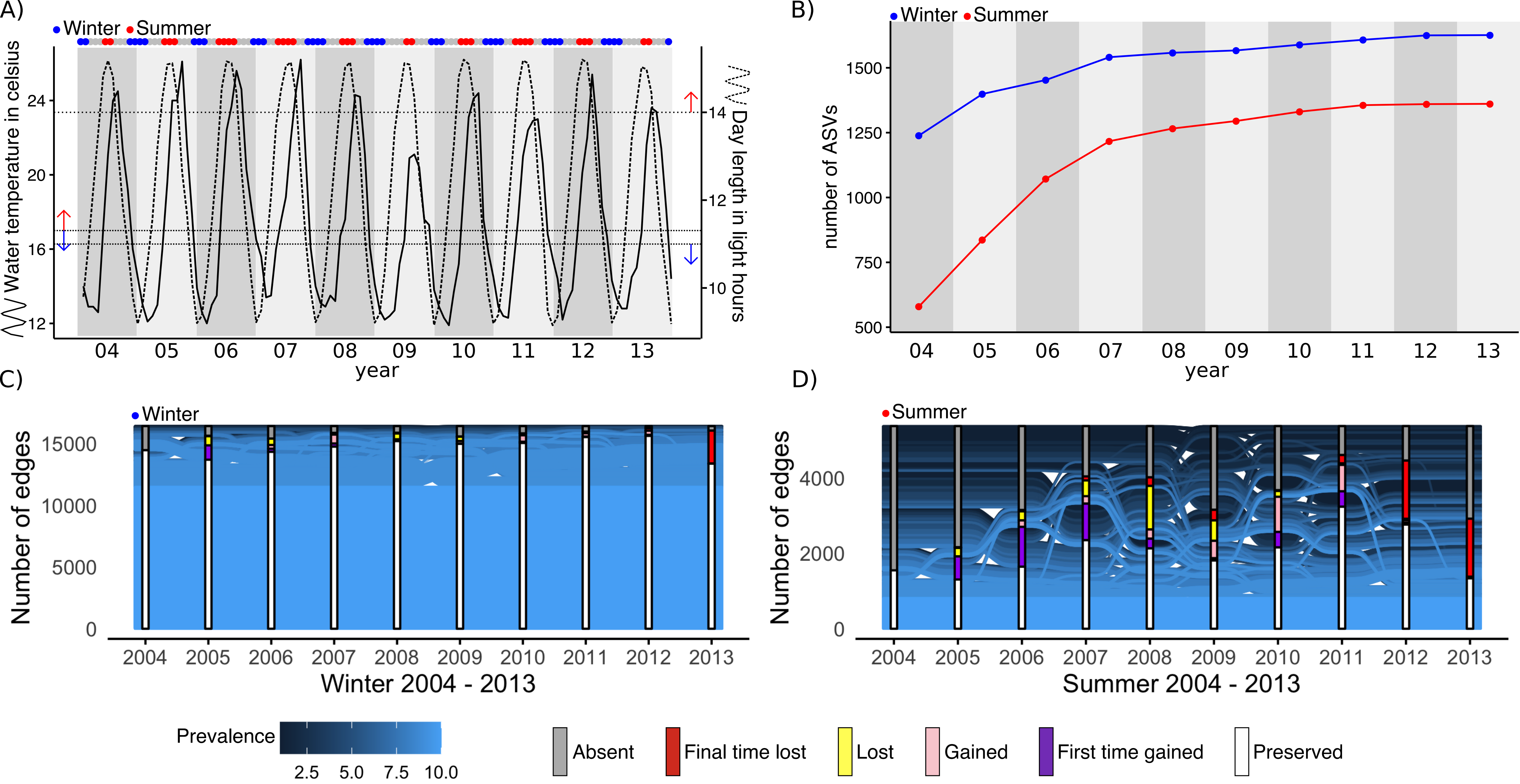

### Supplementary Figure 5

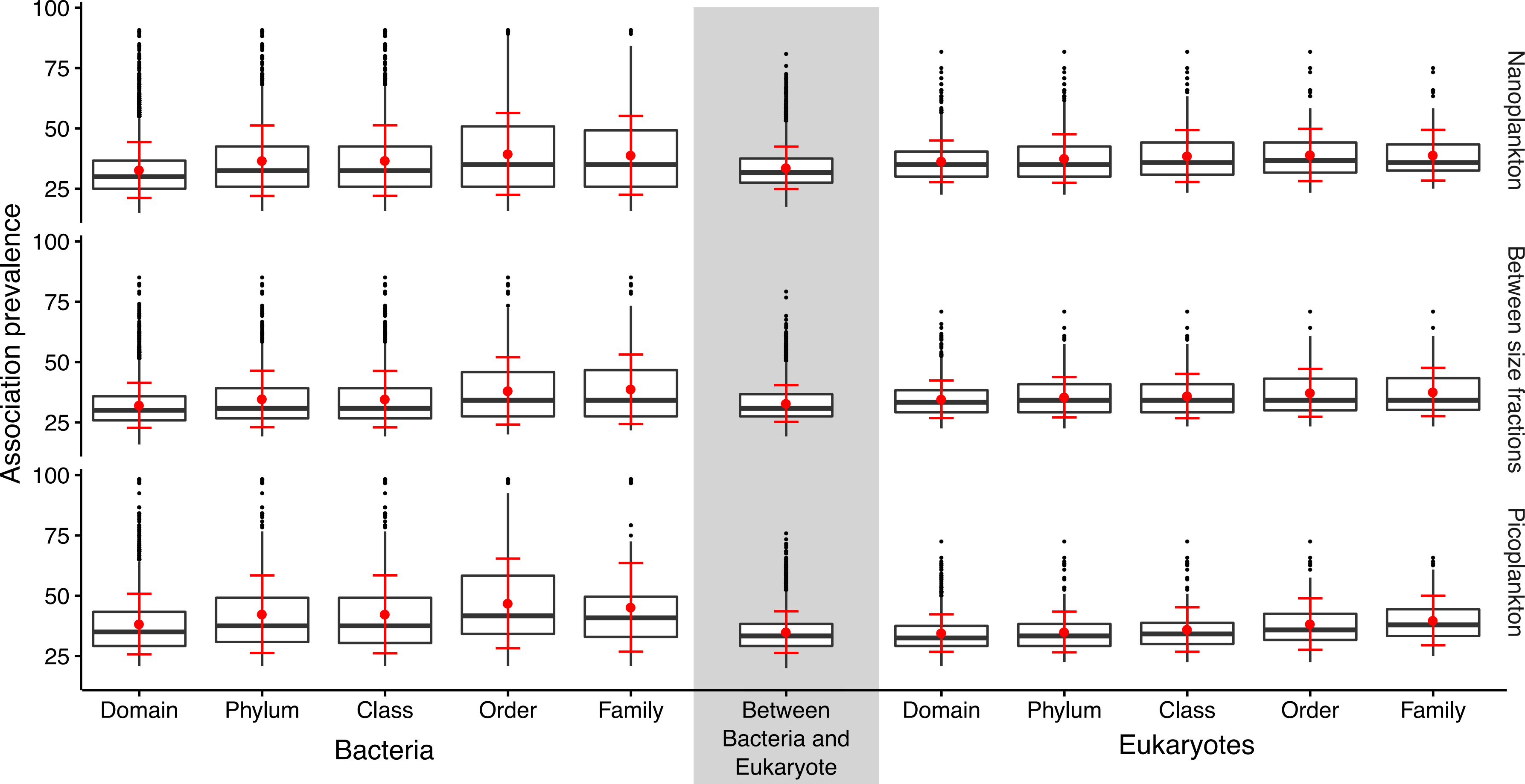

### Supplementary Figure 6

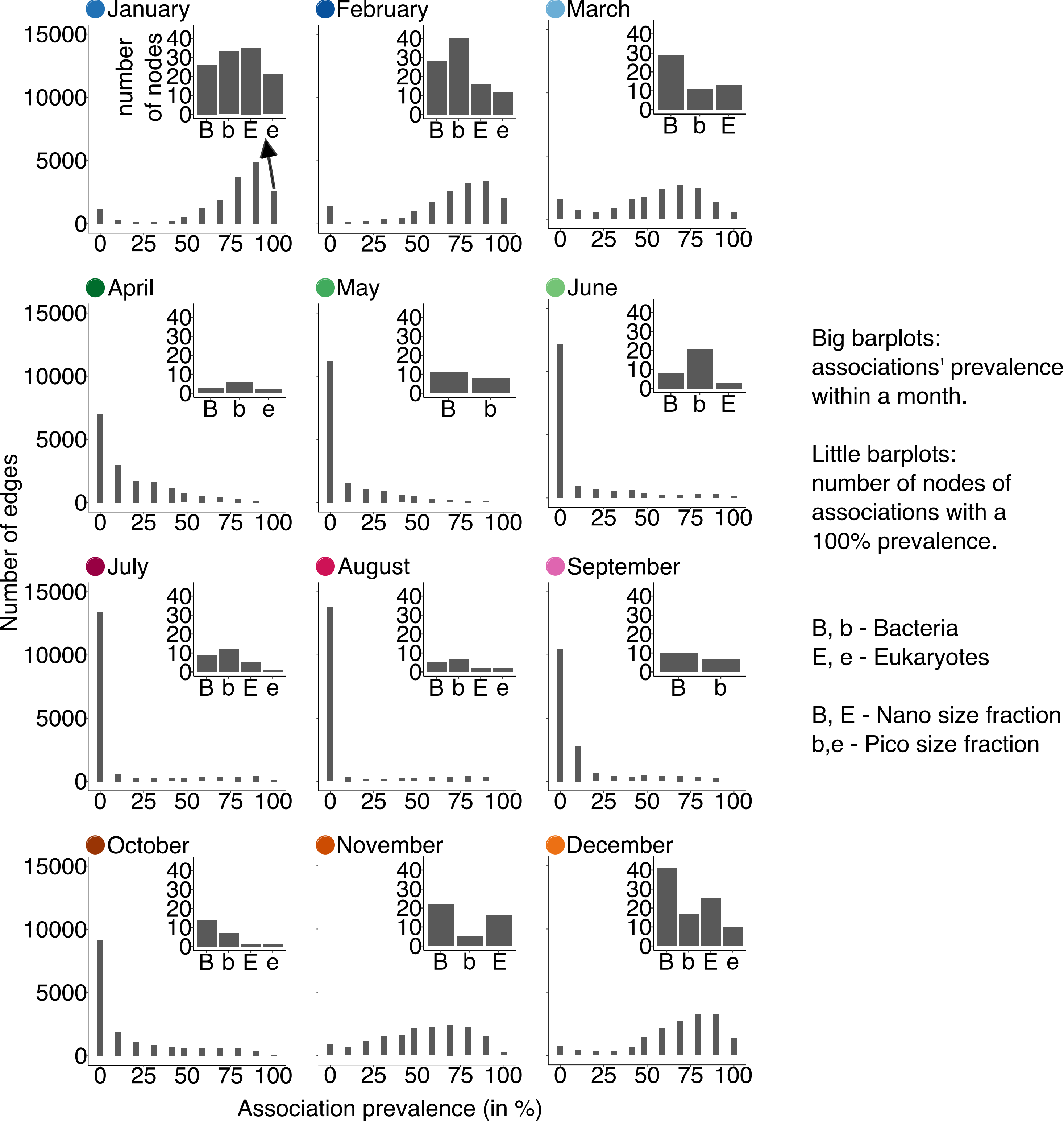

### Supplementary Figure 7

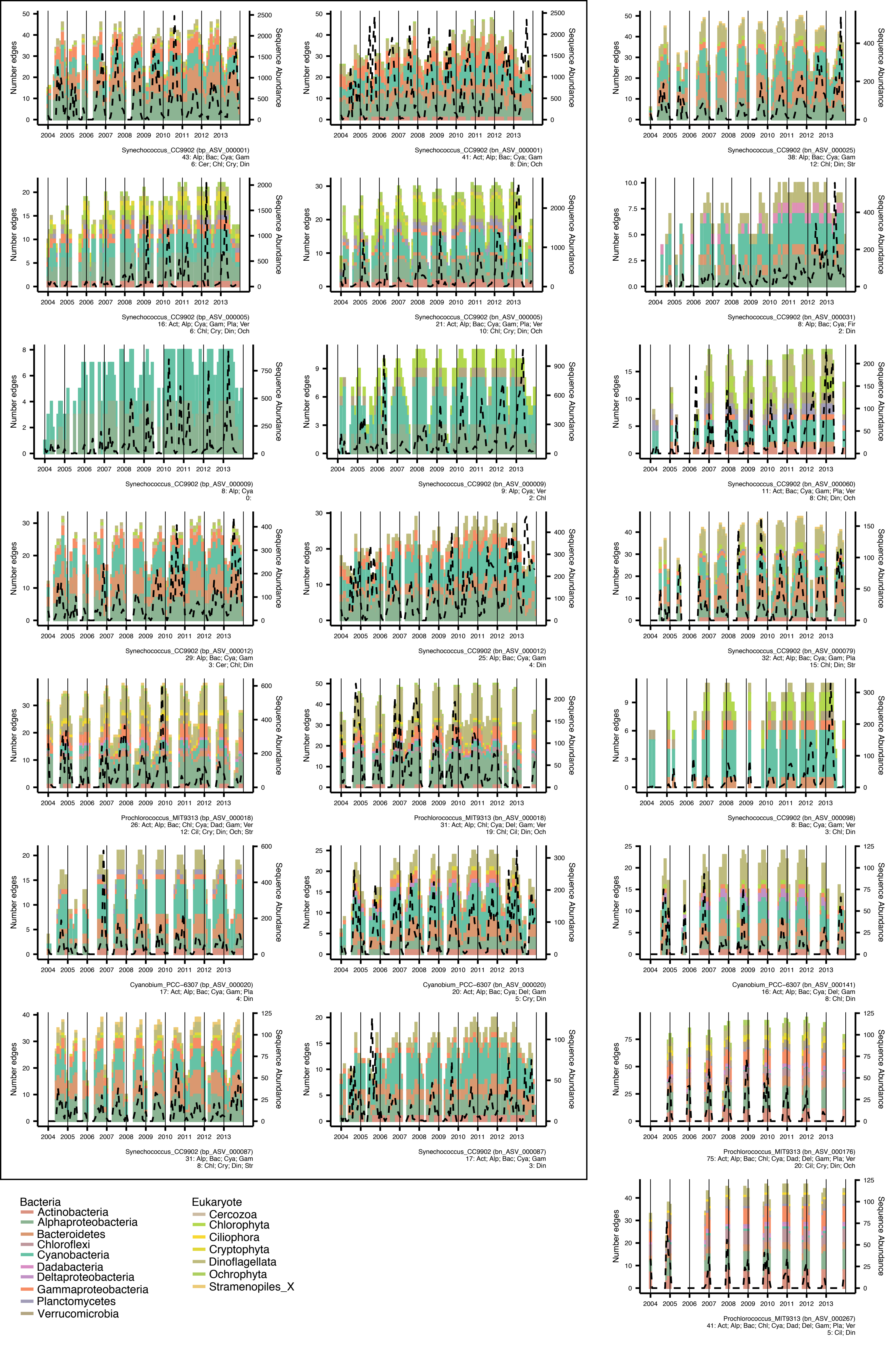

### Supplementary Figure 8

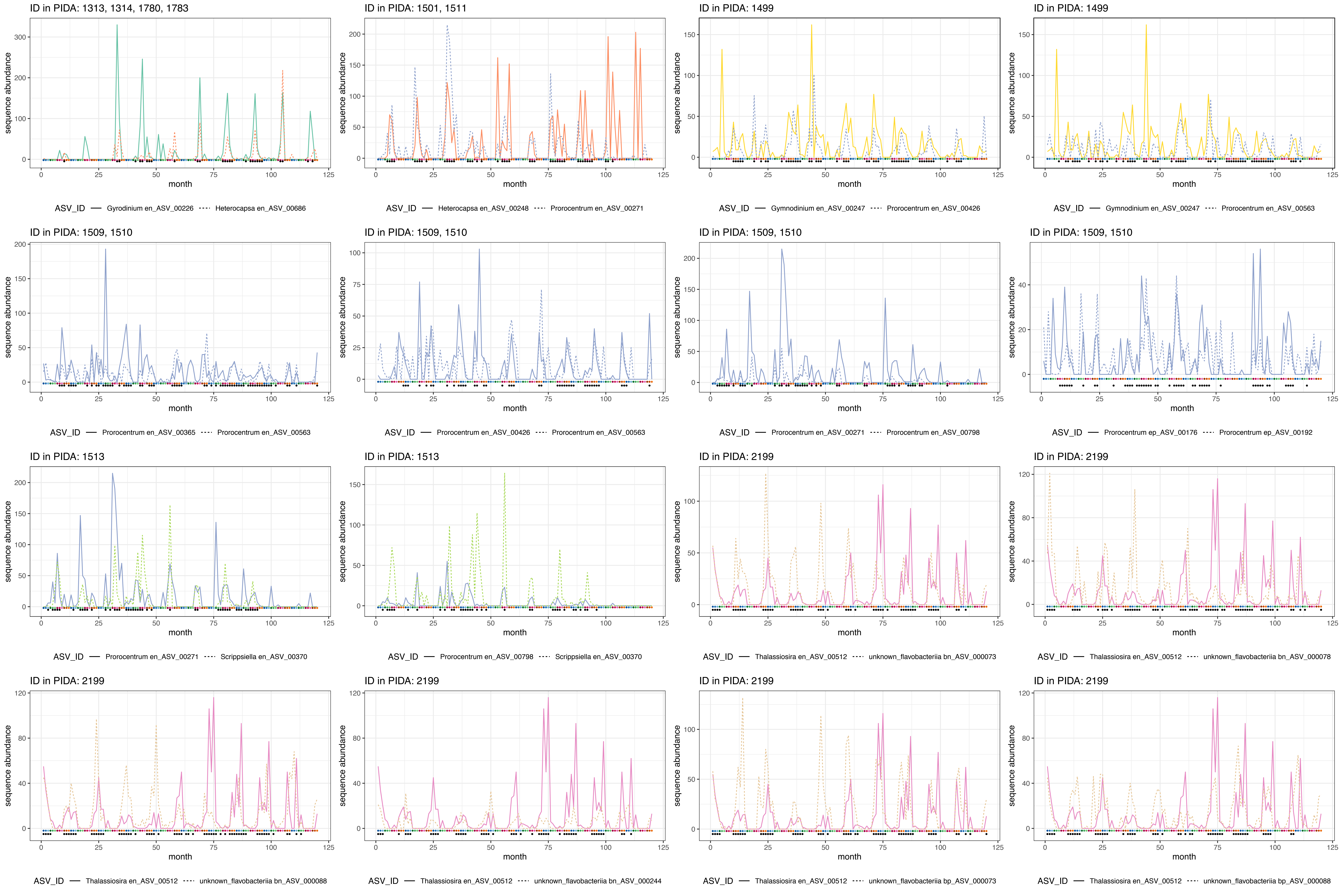
